# The oxytocin system regulates tearing

**DOI:** 10.1101/2022.03.08.483433

**Authors:** Shigeru Nakamura, Toshihiro Imada, Kai Jin, Michiko Shibuya, Hisayo Sakaguchi, Fumiya Izumiseki, Kenji F Tanaka, Masaru Mimura, Kastuhiro Nishimori, Natsumi Kambara, Nozomi Hirayama, Itsuka Kamimura, Kensaku Nomoto, Kazutaka Mogi, Takefumi Kikusui, Yasutaka Mukai, Akihiro Yamanaka, Kazuo Tsubota

## Abstract

Tears are an exocrine physiological fluid secreted onto the ocular surface from the lacrimal apparatus in all mammals. Limited research has been conducted on the functional neuronal circuitry of tear production. In particular, the neuronal mechanisms of emotional tearing, which is a physiological reaction harmonized with enhanced emotional arousal and assumed to be unique to humans, remain unclear. We identified that the oxytocin neurons in the paraventricular hypothalamus is functionally projected to the oxytocin receptor-expressing neurons in the lacrimation center of the superior salivatory nucleus. Optogenetic activation or inhibition of these neurons and/or receptors can modulate the superior salivatory nucleus dependent tear secretion mediated through oxytocin. Moreover, we identified that maternal behavior, nociceptive behavior stimulation, and aversive memory retrieval are linked to tearing in mice, and that these emotional linked tearing are suppressed by optogenetic inhibition of the corresponding oxytocin system. Thus, tearing could be regulated through functional connections between central oxytocin systems in the paraventricular hypothalamus and the superior salivatory nucleus.

## Main text

Tears constitute an exocrine physiological fluid that is secreted onto the ocular surface from the lacrimal apparatus in all mammals via three mechanisms (*1*): 1) basal secretion, comprising a continuous low-level flow that constantly covers the cornea and helps to maintain a healthy transparent ocular surface by ensuring homeostatic balance maintained through basal physiological activity of the lacrimal apparatus; 2) reflex tearing, comprising transient excessive watering from the eye that protects the eye against sudden environmental changes (such as physical and chemical insults) to the ocular surface; and 3) emotional tearing, wherein the evidence for emotional linked tearing in non-human animals is lacking, is triggered by extreme emotional arousal invariably linked to heightened behavioral (vocalizations, kneeling down, jumping around, and/or hugging) and physiological effects (changes in blood pressure, heart rate, respiration, and/or perspiration). Emotional tearing is a salient and expressive visual cue that shapes the perception of human facial expressions and facilitates social bonding among humans (*2*).

Lacrimation is regulated by parasympathetic preganglionic neurons located dorsolateral to the facial nucleus in the dorsolateral portion of the reticular formation, termed the superior salivatory nucleus (SUS). This area sends efferent fibers to the lacrimal gland (LG) via the pterygopalatine ganglion (*3*) and receives afferents from the trigeminal sensory neurons from the ocular surface and various brain regions, including the frontal cortex, basal ganglia, thalamus, and paraventricular hypothalamus (PVH) (*4*)(*5*). Tearing is considered to be regulated by the coordination of these neural systems and LG activity. However, this assumption is based on anatomical findings, largely defined by gross or macroscopic necropsy (*4*), immunostaining (*6*), or non-genetic tracing approaches (*7*), rather than investigations of the functional neural connectivity with cell-subtype specificity.

In particular, for emotional tearing, despite the universality of this phenomenon in humans, the neurological investigation for emotional tearing based on experimental observations has been poorly conducted thus far. The overall purpose of this study was to investigate the functional neuronal circuitry of tearing. Thus, we strived to explore whether mice evoke emotional linked tearing and highlight the neural system that regulates this type of tearing.

## Oxt^PVH^ neurons project to SUS to modulate tear secretion

To shed light on the functional connectivity of the central neural systems that regulate tearing, we initially assessed whether manipulation of SUS parasympathetic neuron activity influences tear secretion. Cre-inducible adeno-associated virus (AAV-FLEX) expressing light-sensitive inhibitory anion-channelrhodopsin-2 (ACR2) fused to mCherry, light-activated cation channel channelrhodopsin-2 (ChR2) fused to enhanced yellow fluorescent protein (eYFP), or mCherry control were unilaterally injected into the SUS of choline acetyltransferase (ChAT)-IRES-Cre mice (*ChAT-Cre* mice), which express Cre recombinase under the control of ChAT promoting the endogenous marker of cholinergic neurons. Fiber optics were implanted above the SUS (Fig. 1A). The histochemical analysis validated the sufficient cell number (Fig. 1B upper bar chart) and co-labeled percentage (Fig. 1B middle bar chart) of the ChAT^sus^ merged with ChR2-eYFP, ACR2-mCherry, or mCherry. No significant differences were noted among ChR2-eYFP, ACR2-mCherry, and mCherry. The ChR2-eYFP, ACR2-mCherry, or mCherry were nearly exclusively co-labeled with ChAT^sus^ (Fig. 1B lower bar chart). Optical stimulation of AAV-FLEX-ChR2-eYFP injected SUS results in significant large c-Fos expression in the ChAT^sus^ neurons co-labeled with ChR2-eYFP compared to that of the optical stimulation of ACR2-2A-mCherry (Fig. S1).

**Figure 1.**
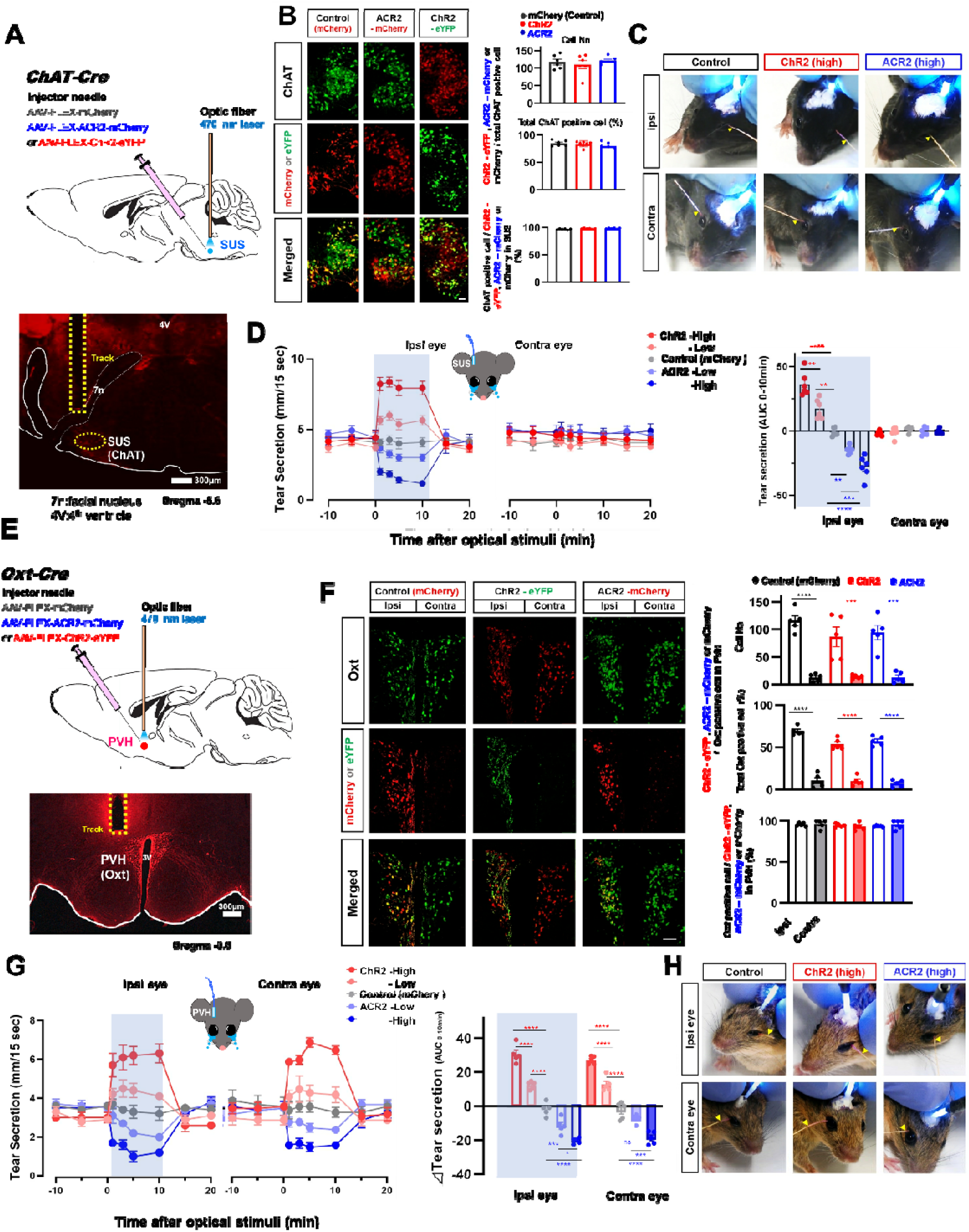
Manipulation of PVH oxytocin neuron activity modifies tear secretion. A. Schematic of optogenetic manipulation of ChAT^SUS^ neurons. Representative image depicting the track of optical fibers above the ipsilateral SUS. B. Histochemical confirmation of the expression of ChR2-eYFP, ACR2-mCherry, and mCherry in the SUS. The right bar chart shows the cell number (upper) and co-localized ratio (middle) of ChAT positive cell merged with ChR2-eYFP, ACR2-mCherry, or mCherry, and co-localized ratio of ChR2-eYFP, ACR2-mCherry, or mCherry with ChAT^SUS^ (lower) in the SUS (n = 5 per group). Scale bar, 20 μm. C. Representative photographs of tear secretion measured with cotton thread during optical stimulation of ChAT^SUS^ neurons. The arrow indicates the wetted length due to tear secretion. D. Dynamics (left) and area under the curve (ΔAUC) _(0-10 min)_ (right) of changes in tear secretion with optogenetic manipulation (n = 5 per group) of ChAT^SUS^ neurons. E. Schematic of optogenetic manipulation of Oxt^PVH^ neurons. Representative image depicting the track of optical fibers above the ipsilateral PVH. F. Histochemical confirmation of the expression of ChR2-eYFP, ACR2-mCherry, and mCherry in the PVH. The right bar chart shows the cell number (upper) and percentage (middle) of ChAT positive cell merged with ChR2-eYFP, ACR2-mCherry or mCherry and co-localized ratio of ChR2-eYFP, ACR2-mCherry, or mCherry with Oxt^PVH^ (lower) in the PVH (n = 5 per group). Scale bar, 100 μm. G. Dynamics (left) and ΔAUC_(0-10 min)_ (right) of changes in tear secretion with optogenetic manipulation (n = 5 per group) of Oxt^PVH^ neurons. H. Representative photographs of tear secretion measured with cotton thread during optical stimulation of Oxt^PVH^→^SUS^ neurons. The arrow indicates the wetted length due to tear secretion. The shaded areas in the figures represent the periods (D left and G left) or side (D right and G right) of optical stimulation. \*\*\*\**P* < 0.0001, \*\*\**P* < 0.001, * *P* < 0.05.

Activation and inhibition of ChAT^SUS^ neurons ipsilaterally increased and decreased tear secretion, respectively. These changes were instantaneous stimulations within the initiation of our measurements (1 min), power-dependent and persistent with optical stimulation. Tear secretion was not altered during optical stimulation in the control group (Fig. 1, C and D and movies S1, A to C).

Oxytocin (Oxt) is a peptide synthesized by neurons predominantly located in the PVH that send descending projections to the SUS (*7*) and supraoptic nucleus (SON), and accessory nuclei (AH) (*8*). Oxt acts as both a hormone and a neuromodulator: the hormonal release of Oxt is essential for parturition and lactation, whereas intracerebral release of Oxt as a neuromodulator has been implicated in the regulation of social *(8, 9)* and emotional processes *(11–13)*. Oxt system-related social and emotional responses that are commonly associated with emotional tearing in humans include fear *(15, 16)*, pain (*9*), distress *(11, 18)*, delight (*10*), and empathy *(20, 21)*. Furthermore, Oxt has been linked to whole-body homeostatic processes (*11*), including food and fluid intake regulation (*12*), thermoregulation (*13*), and cardiovascular regulation (*14*). We therefore functionally interrogated the activity of PVH Oxt (Oxt^PVH^) neurons. AAV-FLEX-ChR2-eYFP, ACR2-2A-mCherry, or mCherry control was unilaterally injected into the PVH of oxytocin-IRES-Cre mice (*Oxt-Cre* mice). Fiber optics were implanted above the PVH (Fig. 1E).

Since the PVH is bilaterally symmetric and lies nearly opposite across the third ventricle, histochemical analysis was performed to validate the ChR2-eYFP, ACR2-mCherry or mCherry expression of contra-ipsilateral PVH. The cell number (Fig. 1F upper bar chart) and colocalized ratio (Fig. 1F middle bar chart) of ipsilateral Oxt^PVH^ merged with ChR2-eYFP, ACR2-mCherry, or mCherry were significantly higher, approximately five fold, than that of the contralateral PVH. No significant differences were noted among ChR2-eYFP, ACR2-mCherry, and mCherry. The ChR2-eYFP, ACR2-mCherry, or mCherry were nearly exclusively colocalized with Oxt^PVH^ (Fig. 1F lower bar chart). Paraventricular hypothalamus c-Fos expression ratio in mCherry-or ChR2-expressing Oxt^PVH^ neurons after optical stimulation are shown in Figure S2. Optical stimulation of AAV-FLEX-ChR2-eYFP injected into the ipsilateral PVH results in the significantly large c-Fos expression in the Oxt^PVH^ neurons merged with ChR2-eYFP in the ipsilateral PVH compared to the contralateral PVH. Upon optical stimulation of AAV-FLEX-mCherry ipsilateral injected into the PVH, no difference was noted in the ipsilateral PVH and contralateral PVH. The expression level in the contralateral PVH upon optical stimulation of AAV-FLEX-ChR2-eYFP was similar to that of mCherry injection (Fig. S2), thereby suggesting that this histochemical validation can be applied to investigate the functional projection from each PVH associated with bilateral eye tear secretion.

Activation and inhibition of Oxt^PVH^ neurons bilaterally increased and decreased tear secretion, respectively. These changes were instantaneous stimulation within the initiation of our measurements (1 min), power-dependent and persistent with optical stimulation similar to that of ChAT^SUS^. Tear secretion was not altered during optic stimulation in the control group (Fig. 1, G and 1H and movies S2, A to C). We further examined the effects of optogenetic activation of the SON or AH on tear secretion. AAV-FLEX-ChR2-eYFP was unilaterally injected into the SON or AH of *Oxt-Cre* mice, and fiber optics were implanted ipsilaterally to the viral injection site. Tear secretion was not altered during optical stimulation of the SON or AH (Fig. S3). These findings resulted in the hypothesis that the interaction between ChAT^SUS^ and Oxt^PVH^ neurons could play a significant role in tearing.

Thereafter, we examined the functional connectivity between Oxt^PVH^ and ChAT^SUS^ neurons. To evaluate whether manipulating the activity of SUS-projecting Oxt^PVH^ (Oxt^PVH^→^SUS^) neurons altered tear secretion, Oxt^PVH^→^SUS^ neurons were modulated by unilateral injections of AAV-FLEX-ChR2-eYFP or AAV-FLEX-ACR2-mCherry into the PVH, and fiber (*15*) optics were bilaterally implanted over the SUS in *Oxt-Cre* mice (Fig. 2A). We observed bilateral representation of the axons expressing ChR2-eYFP or ACR2-mCherry in the SUS region (Fig. 2C). Optical activation and inhibition of the SUS increased and decreased ipsilateral tear secretion from the eye corresponding to the stimulated site, respectively (Fig. 2B). Histochemical analysis of SUS noted that optical stimulation of AAV-FLEX-ChR2-eYFP injected ipsilateral PVH results in significantly large c-Fos expression in the ChAT^sus^ neurons in the bilateral SUS compared to that of optical stimulation of ACR2-2A-mCherry (Fig S5)

**Figure 2.**
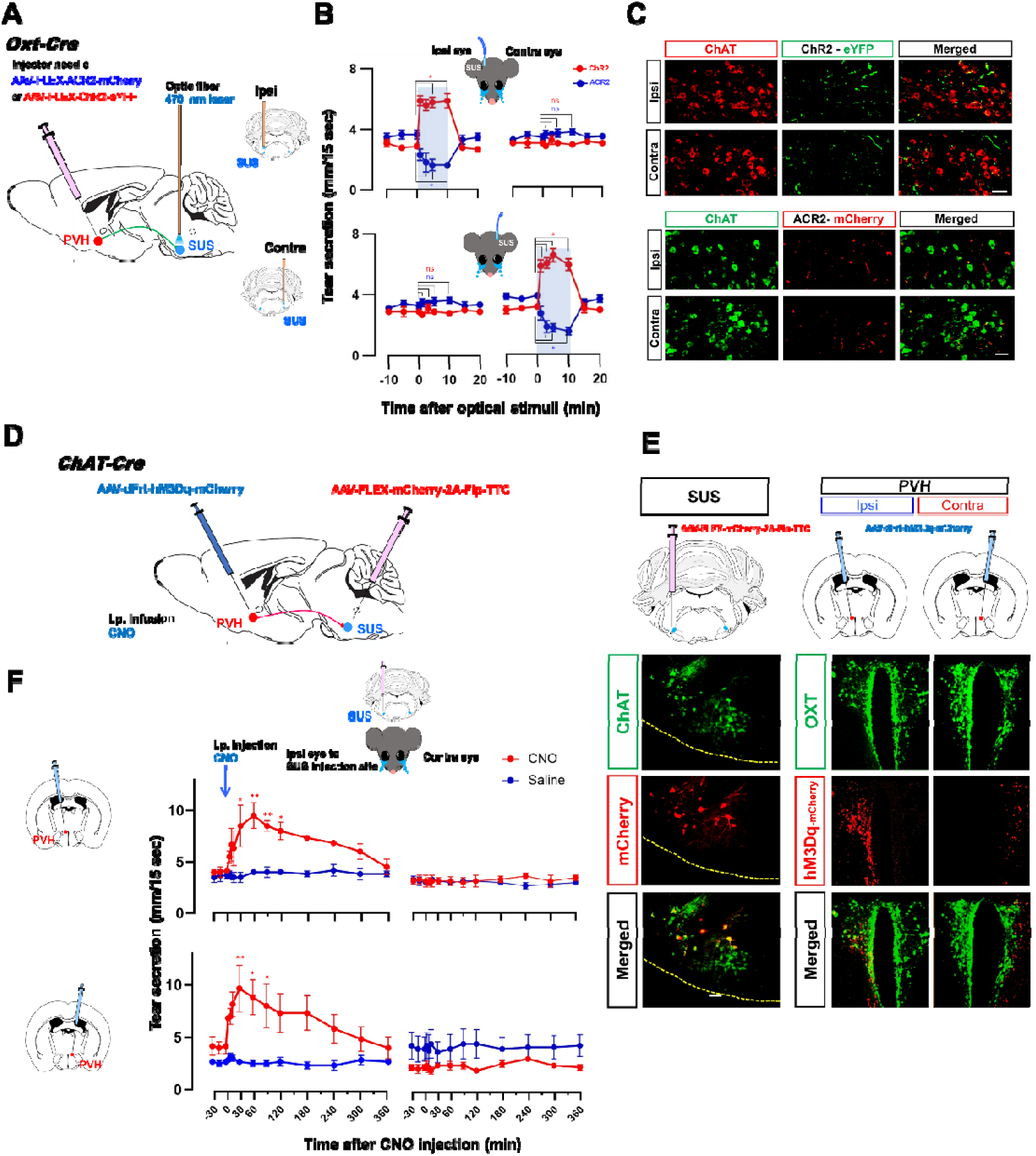
Functional projections of Oxt^PVH^ to ChAT^SUS^ neurons modify tear secretion. A. Schematic of optogenetic manipulation of Oxt^PVH^→^SUS^ neurons. B. Dynamics of tear secretion with optogenetic manipulation of Oxt^PVH^→^SUS^ neurons (n = 4 per group). The shaded areas in the figures represent the periods and side of optical stimulation. \**P* < 0.05 vs 0 min C. Bilateral representation of axons expressing ChR2 -eYFP or ACR2 -mCherry in the SUS region. Scale bar represents 50lµm. D. Schematic of chemogenetic manipulation of PVH neurons projecting onto ChAT^SUS^ neurons using a dual viral vector approach. E. Histochemical confirmation of mCherry expression in the SUS (left) and hM3Dq fused mCherry in the PVH (right). Scale bar represents 50lµm (SUS) and 100 µm (PVH). F. Dynamics of tear secretion with chemogenetic manipulation of PVH neurons projecting onto ChAT^SUS^ neurons (n = 3 per group). \**P* < 0.05, \*\**P* < 0.05 vs 0 min.

A dual viral vector approach was used for chemogenetic activation of the PVH neurons projecting to ChAT^SUS^ neurons (*16*). AAV-dFrt-hM3Dq-mCherry was unilaterally injected into the SUS, and AAV-FLEX-mChrerry-2A-Flippase (Flp)-Tetanus toxin C-fragment (TTC) was injected into the ipsilateral or contralateral PVH corresponding to the virus injected into the SUS of *ChAT-Cre* mice (Fig. 2D). In this system, Flp is trans-synaptically transported to neurons projecting to the ChAT^SUS^ neurons. hM3Dq-expressing AAV was injected into the PVH and expressed DREADDs in a Flp-dependent manner in the ChAT^SUS^-projecting PVH neurons. ChAT^SUS^ injection sites were confirmed by mCherry expression in SUS (Fig. 2E left). We observed the ipsilateral or contralateral expression of hM3Dq fused mCherry in the PVH corresponding to the virus injected into the SUS (Fig. 2E right). Intraperitoneal injection of clozapine N-oxide (CNO) resulted in the unilateral stimulation of tear secretion in the eye corresponding to the SUS viral injection site (Fig. 2F). For tearing, Oxt^PVH^ neuron functionally projects to the bilateral SUS and the effects of each projection from the PVH to SUS on the tear secretory levels is at the same level as the ipsi-or contra-lateral side of the eye. Significant inhibition of the tear response is by the sole ipsilateral inhibition of the Oxt^PVH^→^SUS^ neuron, while still receiving functional stimulation from the contralateral Oxt^PVH^→^SUS^ (Fig. 1G and fig. 2F), implying that the effect of the complicated synaptic transmission, such, as the presence of axoaxonic synapses endings (*17*), may be quite influential on the functional afferent connection at the SUS.

Peripheral Oxt is secreted into the general circulation by Oxt^PVH^ projections to the posterior pituitary gland. A well-established peripheral role of Oxt is milk ejection from the mammary glands during lactation and uterine contraction during parturition via the induction of myoepithelial cell contractions mediated by activation of the peripheral Oxt receptor (OXTR) (*18*). OXTRs are also expressed in the myoepithelial cells of the LG and participate in the excretion of tear fluid, induced by contraction of the LG (*19*). To exclude the possibility that the changes in tear secretion induced by manipulation of Oxt^PVH^ neurons were mediated by peripheral Oxt, a series of optogenetic manipulations in *Oxt-Cre* mice were performed while interrupting neuronal signaling from the central nervous system to the LG via surgical denervation of the lacrimal nerve innervating the LG (*20*). The lobular structure of LG acinar cells was maintained, and an increase in the tear secretion following the application of Oxt or acetylcholine to the LG was confirmed 1day post-surgery suggesting that the LG gland ability is preserved (Fig. S4A). This protocol abolished the increase in tear secretion induced by activation of the Oxt^PVH^ neurons (Fig. S4B). Inhibition of LG OXTRs by exposure of the LG to the OXTR antagonist atosiban did not affect the increase in tear secretion induced by Oxt^PVH^ neuron activation (Fig. S4C). In addition, we observed a significant increase in the c-Fos expression in the ChAT^SUS^ neurons, which was accompanied by Oxt^PVH^ neuron activation (Fig. S5). Collectively, these results demonstrate that functional neuronal connectivity from Oxt^PVH^ to ChAT^SUS^ neurons, rather than peripheral LG OXTRs, regulates the flow of tear secretion.

## SUS expresses OXTR and mediates tear secretion

Given that functional Oxt release from the axonal terminals of Oxt^PVH^ neurons contributes to OXTR-mediated actions (*21*), we examined the expression of OXTR in the lacrimal central SUS. An OXTR-Venus knock-in mouse line, in which the Venus fluorescent protein is expressed under the control of the *OXTR* gene promoter, was used to characterize OXTR-expressing neurons in the SUS region (*22*). We identified a Venus-positive bilateral neuronal population in the SUS region. Approximately 93% of these neurons co-expressed ChAT in *OXTR Venus* (+/-) mice (Fig. 3A). Venus-positive bilateral neuronal populations were not observed in *OXTR Venus* (-/-) mice (Fig. S6).

**Figure 3.**
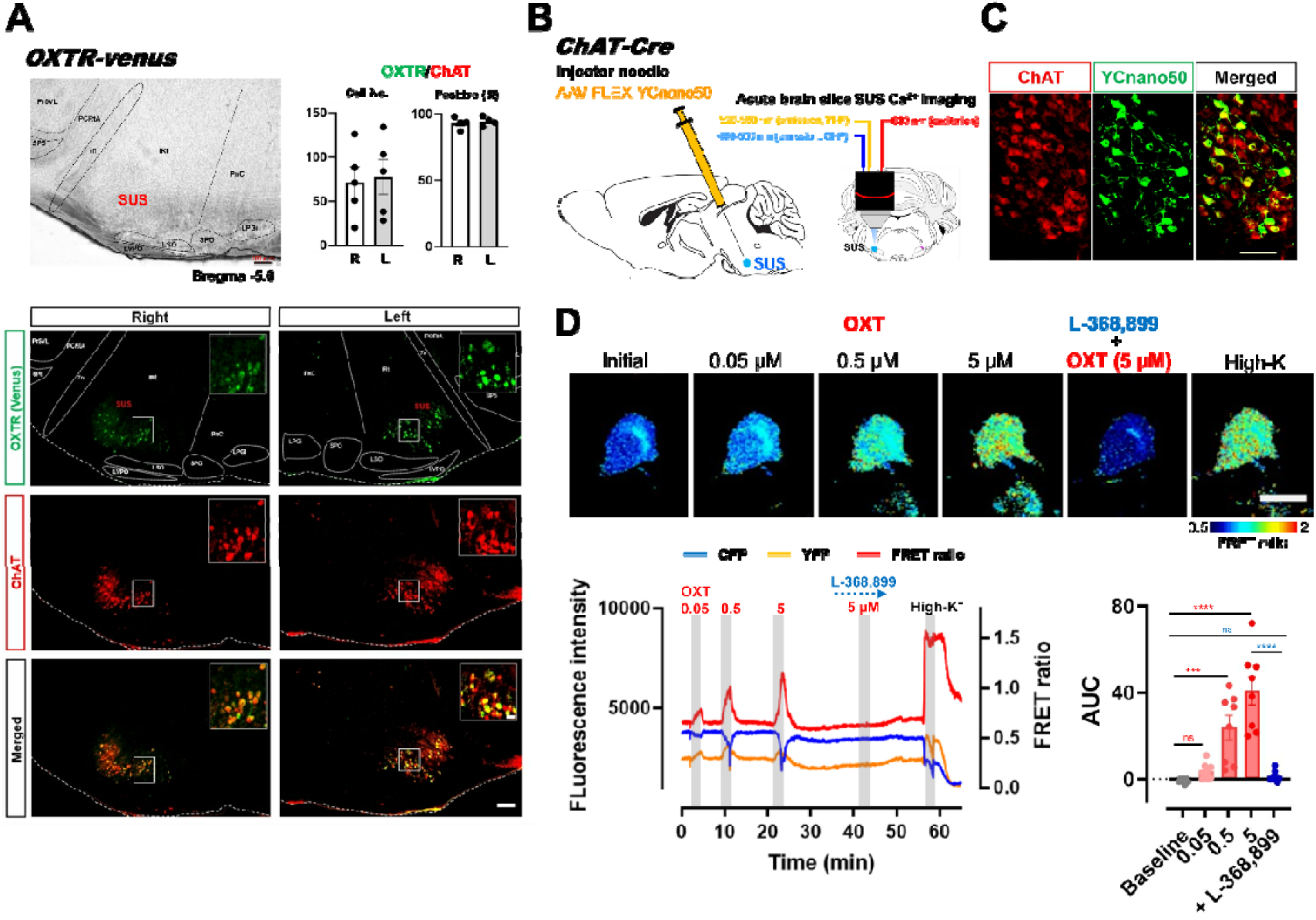
Expression of OXTRs in the SUS. A. Venus fluorescent protein expression in the SUS of *OXTR-Venus* mice (n = 5). Scale bar represents 100lµm or 20 µm (insets). B. Schematic of two-photon Ca^2+^ imaging of ChAT^SUS^ neurons in acute brain slices prepared from *ChAT-Cre* mice. C. YCnano50 expression in the SUS of *ChAT-Cre* mice. Scale bar represents 20lµm. D. FRET ratio changes in ChAT^SUS^ neurons stimulated by Oxt with/without an OXTR antagonist and high K^+^. Pseudo-color images of FRET ratio elevation (upper) and representative changes in fluorescence intensity of cyan fluorescent protein (CFP), YFP, and FRET ratio (lower left). Bar graph indicates the AUC of the increase in FRET ratio (lower right). Scale bar represents 20lµm (n = 8). \*\*\*\**P* < 0.0001, \*\*\**P* < 0.001.

OXTRs activate Gq proteins that couple to downstream signal transduction pathways, thus triggering intracellular calcium responses as secondary messengers to induce various biological reactions (*23*). To confirm whether *OXTR* gene-encoded proteins in ChAT^SUS^ cells exhibited functional properties of endogenous OXTRs, we monitored the activity of ChAT^SUS^ neurons in response to Oxt. To record the activity of ChAT^SUS^ neurons, the Förster resonance energy transfer (FRET)-based ratiometric gene-encoded Ca^2+^ indicator Yellow Cameleon-Nano50 (YCnano50) (*24*) was specifically expressed in ChAT^SUS^ neurons by injecting AAV-FLEX-YCnano50 into the SUS of *ChAT-Cre* mice. We performed two-photon Ca^2+^ imaging of SUS neurons in acute brain slices of the reticular formation including the SUS region (Fig. 3, B and C). Stimulation with Oxt induced a concentration-dependent increase in the FRET ratio in the SUS, and this increase was completely suppressed by application of the selective OXTR antagonist L-368,899 (Fig. 3D and movie S3A). In addition, application of the selective OXTR agonist Way 267,464 increased the FRET ratio in the SUS, and this increase was completely suppressed by L-368,899 (Fig. S7, A to C and movie S3B). Depolarization with high K^+^ induced a transient overshoot in the FRET ratio (Fig. 3D and movie S3, A and B).

To further investigate whether manipulation of OXTR-expressing neurons in the ChAT^SUS^(OXTR^SUS^) modified tear secretion, AAV-FLEX-ChR2-mKate2, ACR2-2A-mCherry, or mCherry control were unilaterally injected into the SUS of OXTR-Cre-GFP mice (*OXTR-Cre* mice). Fiber optics were implanted over the ipsilateral SUS (Fig. 4A).

**Figure 4.**
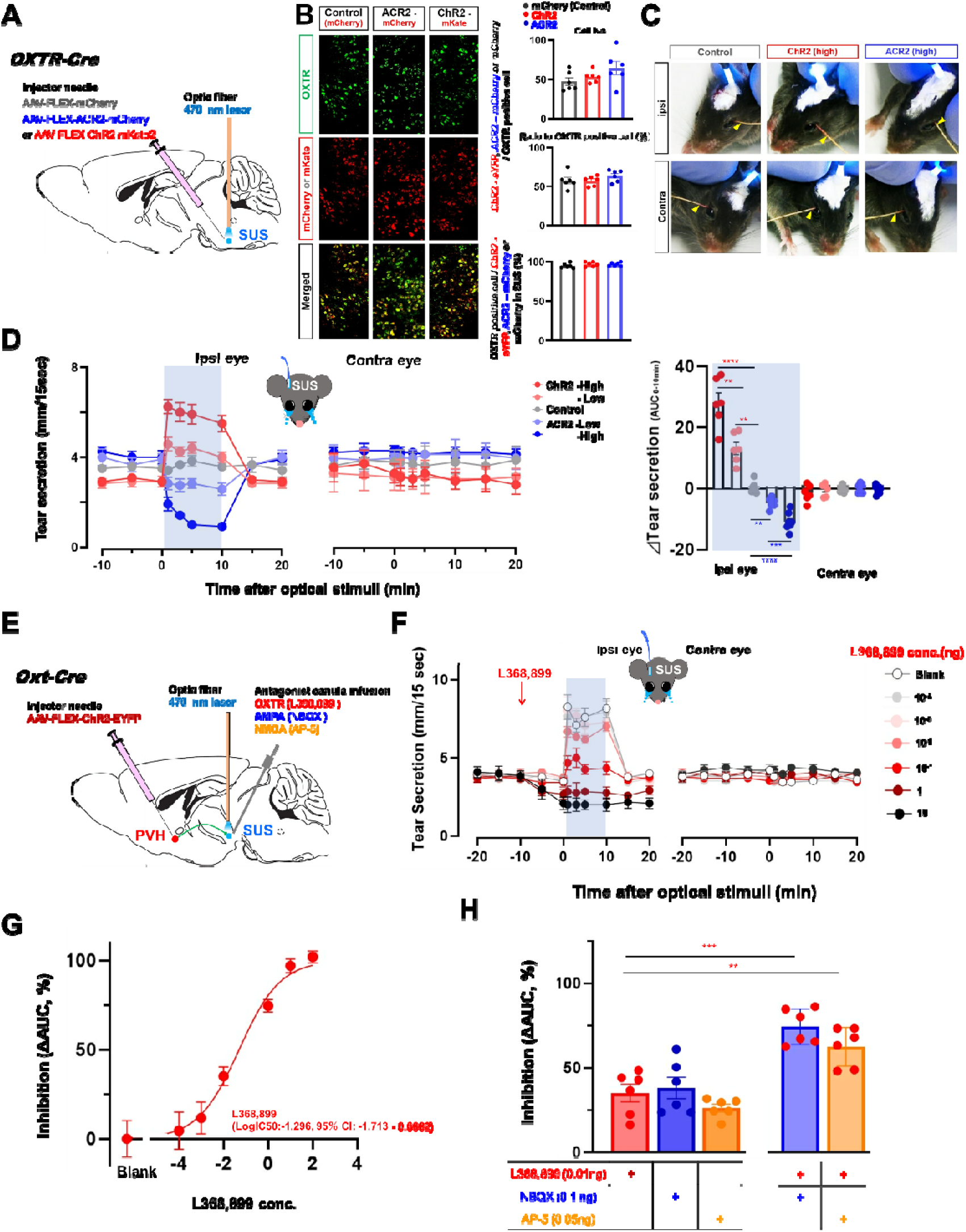
Oxytocin mediates Oxt^PVH^ descending projections to OXTR^SUS^ neurons and modulates tear secretion. A. Schematic of optogenetic manipulation of OXTR^SUS^ neurons. B. Histochemical confirmation of ChR2 fused mKate2, ACR2-mCherry, and mCherry in the SUS. The right bar chart shows the cell number (upper) and co-localized ratio (middle) of OXTR^SUS^ positive cell merged with ChR2 fused mKate2, ACR2-mCherry, or mCherry and co-localized ratio of ChR2 fused mKate2, ACR2-mCherry, or mCherry with OXTR^SUS^ (lower) in the SUS (n = 6 per group). Scale bar represents 50lµm. C. Representative photographs of tear secretion measured with cotton thread during optical stimulation. The arrow indicates the wetted length due to tear secretion. D. Dynamics (left) and ΔAUC_(0-10 min)_ (right) of changes in tear secretion with optogenetic manipulation (n = 6 per group) of OXTR^SUS^ neurons. The shaded area represents the period (left) or side (right) of optical stimulation. E. Schematic of the pharmacological blockade of synaptic transmission of Oxt^PVH^→^SUS^ neurons to downstream neurons in the SUS. F. Dynamics of tear secretion with oxytocin receptor antagonists L368,899 (n = 6 per group). G. Inhibition rate of L368,899. H. Effect of glutamate channel blockers, AP-5 or NBQX alone or co-treatment with L-368,899 on the increase in tear secretion with optical stimulation of Oxt^PVH^→ ^SUS^ neurons. G. The shaded areas in the figures represent periods (D left and F) or side (D right) of optical stimulation. \*\*\*\**P* < 0.0001, \*\*\**P* < 0.001, * *P* < 0.05.

Histochemical analysis validated the sufficient cell number (Fig. 4B upper bar chart) and co-labeled percentage (Fig. 4B middle bar chart) of OXTR^sus^ merged with ChR2 fused mKate2, ACR2-mCherry, or mCherry. No significant differences were noted among ChR2-eYFP, ACR2-mCherry, and mCherry. The ChR2 fused mKate2, ACR2-mCherry, or mCherry were nearly exclusively co-labeled with OXTR^sus^ (Fig. 4B lower bar chart). Optical stimulation of AAV-FLEX-ChR2-mKate2 injected SUS results in significantly large c-Fos expression in the OXTR^sus^ neurons co-labeled with ChR2-eYFP compared to that of optical stimulation of ACR2-2A-mCherry (Fig. S8).

Activation and inhibition of the OXTR^SUS^ neurons ipsilaterally increased and decreased tear secretion, respectively. These changes were stimulation power-dependent and persistent in response to the optical stimulation. Tear secretion was not altered during optical stimulation in the control group (Fig. 4, C and D and movies S4, A toC).

## Oxt partly mediates the link between Oxt^PVH^→^SUS^ and OXTR^SUS^ neurons

Next, to investigate the involvement of Oxt in the increase in tear secretion evoked by the activation of Oxt^PVH^→^SUS^ neurons, we injected AAV-FLEX-ChR2-eYFP unilaterally into the PVH of *Oxt-Cre* mice and implanted an optical fiber and a cannula into the SUS for the pharmacological blockade of synaptic transmission between Oxt^PVH^→ ^SUS^ and downstream neurons. (Fig. 4E). Pretreatment of the SUS with the OXTR antagonist non-peptide L-368,899 dose-dependently decreased basal tear secretion (Fig. S9A) and suppressed the increase in tear secretion induced by optical stimulation of the Oxt^PVH^→^SUS^ neurons (Fig. 4F and 4G). Based on previous reports that Oxt is co-released with glutamate from the axon terminals of the Oxt^PVH^ neurons within the central nucleus of the amygdala (*25*), a series of treatments was performed prior to treating the SUS with glutamate channel blockers. Application of glutamate channel blockers, amino-3-hydroxy-5-methylisoxazole-4-propionic acid (AMPA) receptor antagonist AP-5, or N-methyl-D-aspartate receptor antagonist NBQX alone suppressed the increase in tear secretion induced by the optical stimulation of the Oxt^PVH^→^SUS^ neurons (Fig. 4H, left bar collection). Co-treatment with L-368,899 and AP-5 or NBQX did not compensate for the inhibitory effect of L-368,899 (Fig. 4H, right bar collection). Further, we confirmed that exposure of the SUS to Oxt or OXTR agonists increased tear secretion. Treatment of the SUS with Oxt or the non-peptide OXTR agonist WAY 267,464 via an implanted cannula in WT mice dose-dependently increased tear secretion (Fig. S9, B to D). Suggesting that increased tear secretion modulated by optical stimulation of Oxt^PVH^→ ^SUS^ neurons was modulated via Oxt release within the SUS accompanied by the co-release of another factor that required glutamate channel receptor activation (Fig. S16D).

To further investigate the functional linkage between the SUS-projecting Oxt^PVH^ (Oxt^PVH^→^SUS^) neurons and OXTR^SUS^ neurons in tear secretion, we examined whether an increase in the tear secretion by activation of Oxt^PVH^ is suppressed by synchronized inhibition of the OXTR^SUS^ neurons (Fig. S10). We ipsi-unilaterally injected AAV-FLEX-ChR2-mKate2 into the PVH and AAV-FLEX-hM4Di-mCherry into the SUS of *Oxt*^*Cre/+*^*;OXTR*^*Cre/+*^ mice (*26*), and fiber optics were implanted over the PVH (Figs. S10A). Following intraperitoneal injection of CNO, ipsilateral sustained decrease in the basal tear secretion was observed (Fig. S10C gray line). While basal tear secretion remained unchanged, bilateral increase in tear secretion during optical stimulation of the PVH was observed (Fig. S10C, black line) upon saline pre-injections with optical stimulation of the PVH. For the CNO pre-injections with optical stimulation of the PVH, under decrease in the basal tear secretion, ipsilateral suppression of increased tear secretion during optical stimulation of PVH was observed (Fig. S10C, red line). Histochemical analysis confirmed the bilateral expression of ChR2 fused mKate2 and ipsilateral expression of hM4Di fused mCherry in the OXTR^SUS^ (Fig. S10B). To further ensure that the mCherry-positive neurons in the SUS of *Oxt*^*Cre/+*^*;OXTR*^*Cre/+*^ mice originated from the OXTR^SUS^ and not the Oxt^PVH^ neurons, we injected AAV-FLEX-ChR2-mKate2 into the PVH of *OXTR*^*Cre/+*^ mice or injected the AAV-FLEX-hM4Di-mCherry into the SUS of the *Oxt*^*Cre/+*^ mice. During optical stimulation of PVH of the *OXTR*^*Cre/+*^ mice did not alter the tear secretion, and no visible expression of ChR2 fused mKate2 was observed in the OXT^PVH^ (Fig. S10D). Intraperitoneal injection of CNO into the *Oxt*^*Cre/+*^ mice did not alter the tear secretion, and no visible expression of hM4Di fused mCherry was observed in the ChAT^SUS^ (Fig. S10E).

Collectively, these data highlighted that neuronal signaling from Oxt^PVH^ neurons to OXTR^SUS^ neurons mediated via Oxt released from Oxt^PVH^ neurons at least partly mediates tear secretion originating from the central nerve system

## Oxt^PVH^→^SUS^ neurons are involved in basal but not reflex tearing

Tearing is categorized into three types based on their roles: basal, reflex, and emotional (*1*), we examined whether Oxt^PVH^→^SUS^ and/or OXTR^SUS^ participates in tearing according to these categories.

Optogenetic activation and inhibition of Oxt^PVH^→^SUS^ and/or OXTR^SUS^ neurons increased and decreased basal tear secretion, respectively (Fig 1G, fig. 2A and fig. 4D). These Oxt-related neurons may be involved in the maintenance of sustained basal tearing for providing a wettable and healthy transparent ocular surface (Fig. S16A).

Reflex tearing constitutes transient tearing evoked by sensory stimuli to the ocular surface, such as cold sensation, foreign bodies, or irritant chemicals (*27*). Reflexive tears are secreted regardless of the mental or emotional status. Sensory stimuli received by sensory receptors on the ocular surface are afferently transmitted to the SUS via the ophthalmic branch of the trigeminal nerve and re-transmitted to the efferent pathways, which reach the LG and stimulate tear secretion (*28*). To examine the involvement of Oxt^PVH^→ ^SUS^ and OXTR^SUS^ neurons in reflex tearing, we measured the changes in the secretory capacity of reflex tearing under the optogenetic inhibition of Oxt^PVH^→^SUS^ or OXTR^SUS^ neurons. To inhibit Oxt^PVH^→^SUS^ neurons, we bilaterally injected AAV-FLEX-ACR2-mCherry into the PVH and implanted fiber optics over the uilateral SUS of *Oxt-Cre* mice (Fig. 5A). To inhibit OXTR^SUS^ neurons, we injected AAV-FLEX-ACR2-mCherry into the unilateral SUS and implanted fiber optics over the ipsilateral SUS of *OXTR-Cre* mice (Fig. 5C).

**Figure 5.**
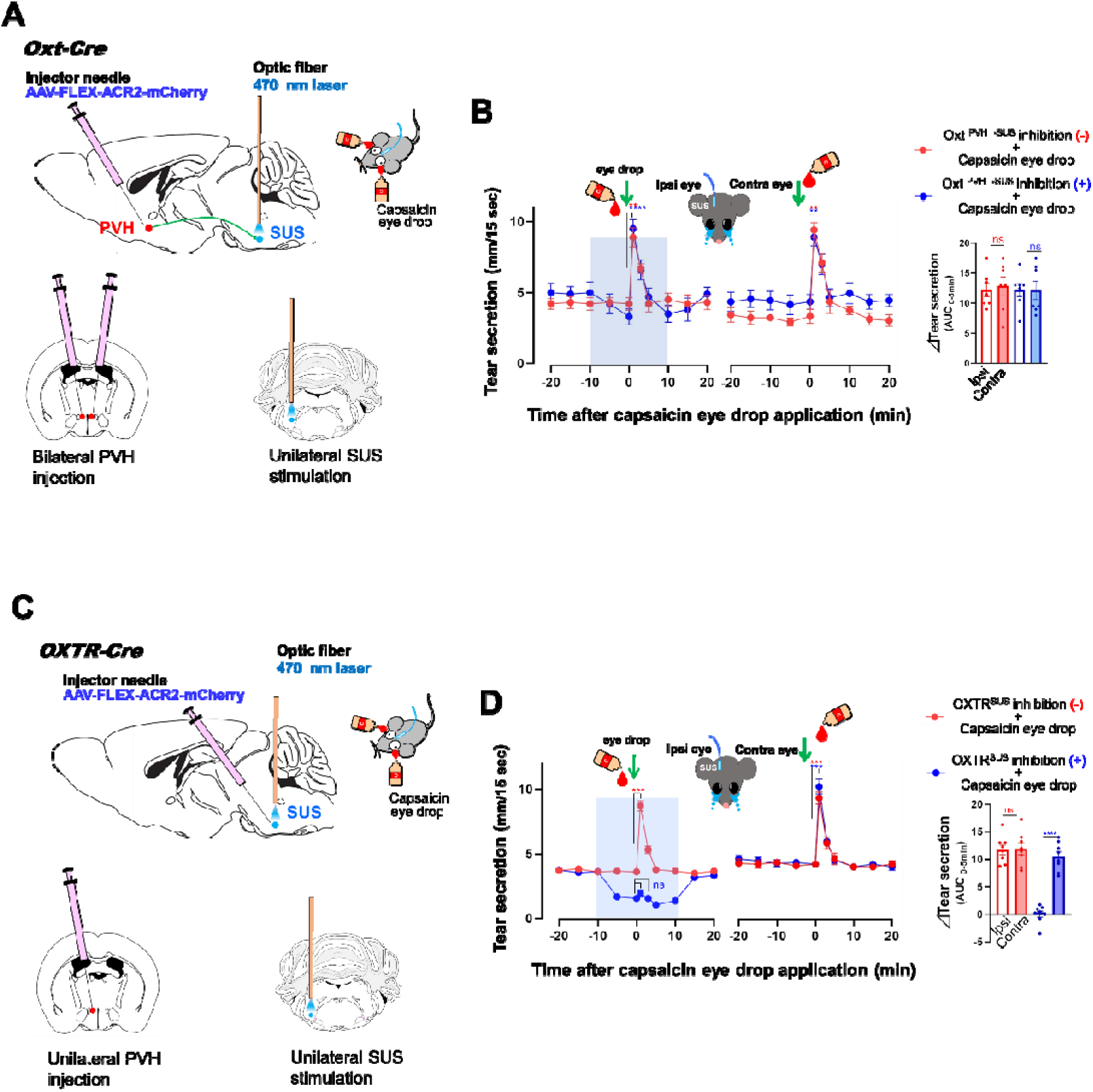
Oxt^PVH^→^SUS^ neurons are not involved in reflex tearing. A, B. Effect of optogenetic silencing of Oxt^PVH^→^SUS^ neurons on reflex tearing (n = 7 per group). C, D. Effect of optogenetic silencing of OXTR^SUS^ neurons on reflex tearing (n = 8 per group). H. The shaded areas in Panels B and D represent the period of optical stimulation. \*\*\*\**P* < 0.0001, \*\*\**P* < 0.001, \*\**P* < 0.01, * *P* < 0.05.

Transient reflex tearing was evoked by sensory eye stimulation with topical eye instillation of an agonist of nociceptor transient receptor potential vanilloid 1, capsaicin (*29*). Following bilateral inhibition of Oxt^PVH^ neurons projection to unilateral SUS (Fig. 5A), the capsaicin-stimulated transient increase in tear secretion did not change with or without the inhibition of Oxt^PVH^→^SUS^ neurons (Fig. 5B). However, unilateral inhibition of OXTR^SUS^ neurons (Fig. 5C) suppressed capsaicin-stimulated tear secretion from the eye ipsilateral to inhibited OXTR^SUS^ neurons (Fig. 5D). Instillation of vehicle eye drops did not alter tear secretion with or without the inhibition of Oxt^PVH^→ ^SUS^ (Fig. S11A)or OXTR ^SUS^ neurons (Fig. S11B), suggesting that OXTR^SUS^ neurons and not Oxt^PVH^→^SUS^ neurons participate in reflex tearing (Fig. S16B).

## Emotional-linked tearing in mice

Emotional tearing with bilateral synchronization is associated with increased tear secretion accompanied by high emotional arousal. Rodent models are used extensively to elucidate the neural mechanisms of the sleep-wake cycle, memory system, social cognition, and sensory transduction. However, there is a paucity of research on emotional tearing in rodents, as this phenomenon is assumed to be unique to humans despite lack of conclusive evidence.

The term “Emotion” can be referred to as an adaptative behavioral response that allows an organism to increase its chance for survival and can be traced back to at least Darwin (1872) (29). This phenomenon related to survival functions includes defense, maintenance of energy and nutritional supplies, fluid balance, thermoregulation, and reproduction. Likewise, it is proposed that animal emotions are evolved from basic mechanisms that were derived from positive (seeking, lust, care, play) and negative (fear, rage/anger and sadness/panic) responses (*30*). The neural systems that modulate these survival functions regard emotion to be conserved to a significant degree across mammalian species, including humans (*31*). Thus, in order to assess potential emotional-like tearing in mice, we measured the increase in bilateral tear secretion under various behavioral stimulatory conditions based on the emotions with positive (bonding and reward) or negative (nociception and distress) valence. Among the behavioral stimulation conditions, suckling dams (a behavior associated with maternal bonding) and persistent nociception induced by paw formalin injections (*32*) and facial formalin injections (*33*) elicited marked increases in bilateral tear secretion (Fig. 6A). While the response to persistent nociception or maternal stimuli that increased tearing has significant importance for survival function, the common features among behavioral stimuli in the presence or absence of tearing (Fig. 6A) are unclear. Considering that the expression of the survival-related behaviors is species-specific (*31*), the observed variance among tearing in response to behavioral stimuli may possibly result from the differences in the relative importance for their own survival in mice.

**Figure 6.**
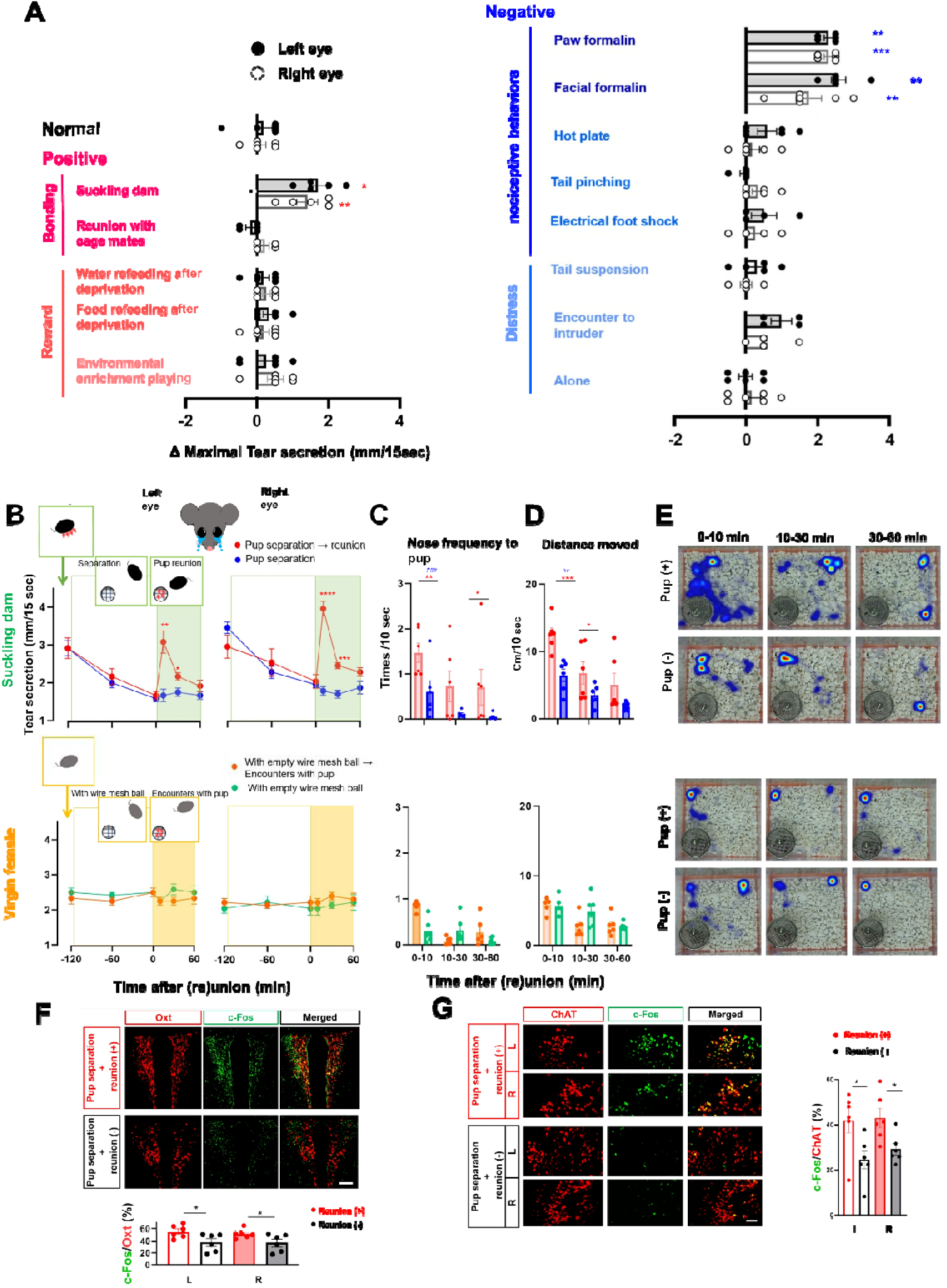
Mouse maternal behavior-linked tearing. A. Effects of various behavioral stimulations based on tear secretion (n = 6 per group with the exception of n = 4 per group for electric foot shock and Encounter to intruder, n=5 per group for Suckling dam, Paw formalin and Tail suspension). \**P* < 0.01, \*\**P* < 0.01 vs NT eyes. B. Changes in tear secretion during mother and pup separation and reunion in dams (upper) and virgins (lower). \*\**P* < 0.01, \**P* < 0.05 vs pup separation with reunion (-). C−E. Mouse pup retrieval behavior. Frequency of nose contact with pups (C), distance moved (D), and heat map of tracking (E). \*\**P* < 0.01, \**P* < 0.05 vs pup separation with reunion (-), ## *P* < 0.01, ###*P* < 0.05 vs virgin encountered with pup. I. F, G. c-Fos expression in the PVH (F) region and SUS (G). The bar chart shows percentage of the Oxt ^PVH^ (F) and ChAT ^SUS^ (G) cells merged with c-Fos (n = 6 per group). Scale bar represents 100 µm and 50lµm, respectively. \**P* < 0.05 vs reunion (-).

To further investigate whether the increase in tearing in response to behavioral stimulation conditions could be regarded as emotion-linked tearing, we focused on suckling dams and paw formalin injections, the latter being a well-established protocol for nociceptive behavior (*32*). Maternal behavior comprises a highly conserved motivated behavioral repertoire in mammals that is indispensable for the survival and development of pups (*34*). To characterize tearing elicited in suckling dams during maternal bonding, we evaluated the changes in tear secretion in suckling dams during mother pup separation and reunion. Pups were encased in a wire mesh ball and were reunited with dams following 2 hours of separation in independent housing to avoid social interaction. In suckling dams, tear secretion decreased to normal values during pup separation. After reunion, tear secretion temporarily returned to the same levels as those during the suckling period at 10 min and subsequently decreased to the same levels as those during the separation period (Fig. 6B, red line, upper panel). In suckling dams that were not reunited with pups, tear secretion did not change during an encounter with an empty wire mesh ball (Fig. 6B, blue line, upper panel). Pup retrieval behavior, frequency of nose contact with pups (Fig. 6, C and E, upper panel), and distance moved (Fig. 6,D and E, upper panel) were significantly increased during the first 10 min after reunion with pups and remained at the same levels thereafter compared to the levels in the dams that were not reunited with pups. Further, we examined c-Fos expression in the PVH and SUS. c-Fos expression in the PVH (Fig. 6F) and SUS (Fig. 6G) was higher in dams that were reunited with pups than in the dams that were not reunited with pups. In virgin females, tear secretion was not changed by encounters with pups or non-pup stimuli, and basal values were sustained throughout the evaluation period (Fig. 6B, lower panel). The frequency of nose contact with pups was significantly higher in the first 10 minutes after encountering pups compared to that without pups, although the values were significantly lower than those for suckling dams (Fig. 6, C and E, lower panel). Distance moved did not change after encountering pups or non-pup stimuli during the monitoring period (Fig. 6, D and E, lower panel).

Although the precise emotions and feelings associated with crying may vary, human crying is generally elicited by emotional arousal from a combination of external stimuli (e.g., social situations and physical stimulation) and/or internal stimuli (e.g., thoughts, memories, and experiences) (*35*). To investigate whether mouse tearing could be linked to internal-external emotional cues, we evaluated the relationship between tearing and contextual conditioning of aversive memories using paw formalin injections. Paw formalin injections were performed in the context condition, and changes in the tear secretion alongside nociception responses were measured. Retrieval and extinction of aversive memories were examined in the same context condition without formalin treatment, and changes in tear secretion were measured at intervals of 1 to 3 days (Fig. 7A).

**Figure 7.**
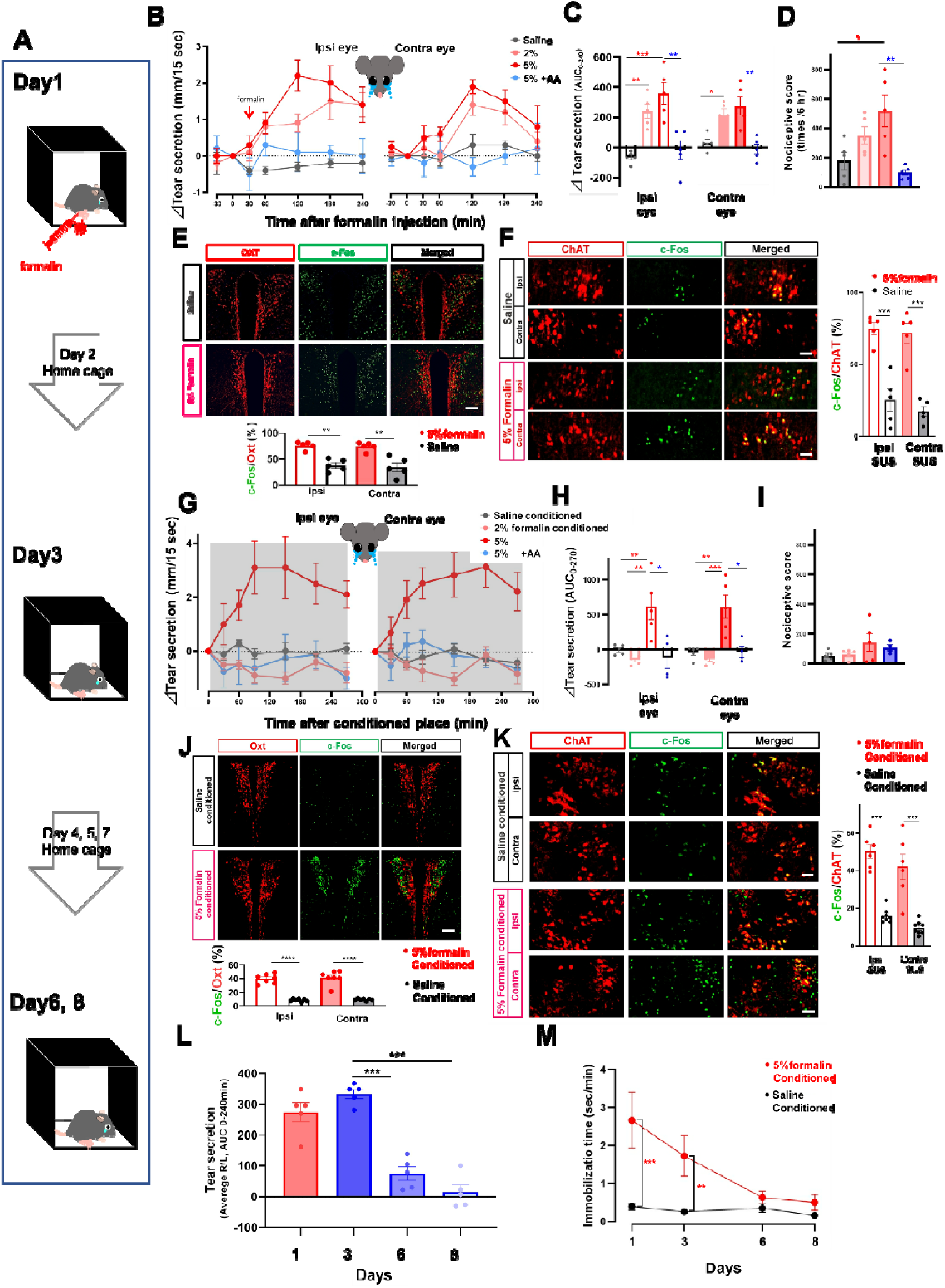
Mouse nociceptive behavior and contextual conditioning of aversive memory-linked tearing. A. Experimental schedule of nociceptive behavior stimulation and aversive memory-induced tearing following paw formalin injections. B−E. Relationship between tearing and nociceptive responses in PF in the context condition (day 1). Effect of formalin dosage on tear secretion dynamics (B) and ΔAUC _(0-10 min)._ (C) Nociceptive responses (D) (n = 5 per group). E−F. c-Fos expression in the PVH (E) and SUS (F) regions after PF (n = 6 per group). Scale bars represent 100 µm and 50lµm, respectively (F). G−I. Reexposure to the context condition on the day after formalin injection (day 3). Tear secretion dynamics (G) and ΔAUC _(0-10 min)._ (H), nociception response (I). The shaded area represents the periods of reexposure to the context condition (G). J−K. c-Fos expression in the PVH (J) and SUS (K) region on day 3. Scale bars represent 100 µm and 50lµm, respectively (J). L−M. Changes in tearing (L) and immobilization time (M) from days 1 to 8 (n = 5 per group). \*\*\*\**P* < 0.0001, \*\*\**P* < 0.001, \*\**P* < 0.01, * *P* < 0.05.

An increase in tear secretion was associated with the dose of formalin injections and nociceptive responses (Fig. 7, B to D) after paw formalin injections (day 1). Suppression of nociceptive responses by pre-treatment with the nonsteroidal analgesic acetylsalicylic acid (AA) diminished the increase in tear secretion (Fig. 7, B to D). In addition, paw formalin injections increased tear secretion, and these effects were attenuated after surgical denervation of the lacrimal nerve innervating the LG (Fig. S12) (*20*). This demonstrated that the increase in tearing was not due to peripheral stimulation of the LG by inflammatory factors caused by systemic inflammatory responses.

During re-exposure to the context condition on the day after formalin injection (day 3), the changes in the tear secretion patterns were similar, based on the dose of the formalin injections and suppression with AA, to those of day 1 (Fig. 7, G and H). Nociceptive responses were not observed during re-exposure in the absence of paw formalin injections (Fig. 7I). On days 6 and 8, tear secretion did not increase upon re-exposure (Fig. 7L). Immobilization time of the 5% formalin conditioned group was significantly longer on days 1 and 3 compared to that of the saline conditioned group (Fig. 7M). We also examined c-Fos expression in the SUS and PVH on days 1 and 3. Following treatment with formalin, c-Fos expression in the SUS (Fig. 7E) and PVH (Fig. 7F) was higher than that following saline injections.

To further confirm whether the increase in tear secretion was associated with the context condition, changes in tear secretion during retrieval and re-exposure were compared among three groups: (i) home cage chamber with bedding (HC), a two-chambered preference apparatus in which the context condition box was connected with (ii) the same context chamber (CC-CC), and (iii) the home cage chamber with bedding (CC-HC) separated by a removable guillotine door to confine the animal (Fig. S13A). The separation door was opened 30 min after re-exposure to allow the mouse to passively cross the apparatus. Tear secretion did not change in the HC group. In the CC-CC group, the changes in the tear secretory patterns were similar to those during re-exposure to the context condition, and no apparent effect of opening the separation door was noted. In the CC-HC group, the increase in tear secretion induced by the context condition decreased at the time of opening of the separation door and returned to its initial value within 180 min (Fig. S12C). Preference among two chambers after opening of the separation door was equivalent in the CC-CC and biased toward the HC camber in the CC-HC. (Fig. S13B). Contextual fear conditioning by electrical foot shock, nociceptive behavior stimulation that did not evoke tearing (Fig. 6A) did not alter the tear secretion. This may imply that distinctive features and/or intensity of emotional stimuli are necessary for evoking a tearing response (Fig. S14).

Finally, we evaluated the involvement of Oxt^PVH^→^SUS^ neurons in elicited tearing during mother pup separation and reunion (Fig. 8B), nociceptive behavior stimulation (Fig. 8C), and aversive memory retrieval (Fig. 8D) following paw formalin injections. Changes in tear secretion were measured under the inhibition of Oxt^PVH^→^SUS^ neurons in *Oxt-Cre* mice. Mice were unilaterally injected with AAV-FLEX-ACR2-mCherry or AAV-FLEX-mCherry control into the PVH, and fiber optics were implanted over the ipsilateral SUS (Fig. 8A). Under the mother pup separation and reunion, optical stimulation was performed during 30minutes before to 10 minutes after reunion. Increase in bilateral tear secretion by pup reunion was evoked in mCherry with/without and ACR2 without optical stimulation (Fig. 8B). Unilateral inhibition of Oxt^PVH^→^SUS^ neurons by ACR2 with optical stimulation suppressed tearing from the ipsilateral eye (Fig. 8B). Pup retrieval behavior, frequency of nose contact with pups (Fig. S15A), and distance moved (Fig. S15B) were as same level at mCherry and ACR with/without optical stimulation. In the nociceptive behavior stimulation and following aversive memory retrieval, optical stimulation was performed during 60 minutes after paw formalin injection or re-exposure to the context condition. Increased in bilateral tearing was evoked by paw formalin injection (Fig. 8C right) and re-exposure to the context condition (Fig. 8D right) was as same level at ACR2 and mCherry. During the optical stimulation, increase in tearing from ipsilateral eye was significantly suppressed with inhibition of Oxt^PVH^→^SUS^ in ACR2 compare to mCherry (Fig. 8, C and D). Overall, our results suggest that emotionally linked tearing during maternal behavior, nociceptive behavior stimulation, and aversive memory retrieval is triggered via activation of Oxt^PVH^ neurons that project to the lacrimation central SUS in response to emotional arousal (Fig. S16C).

**Figure 8.**
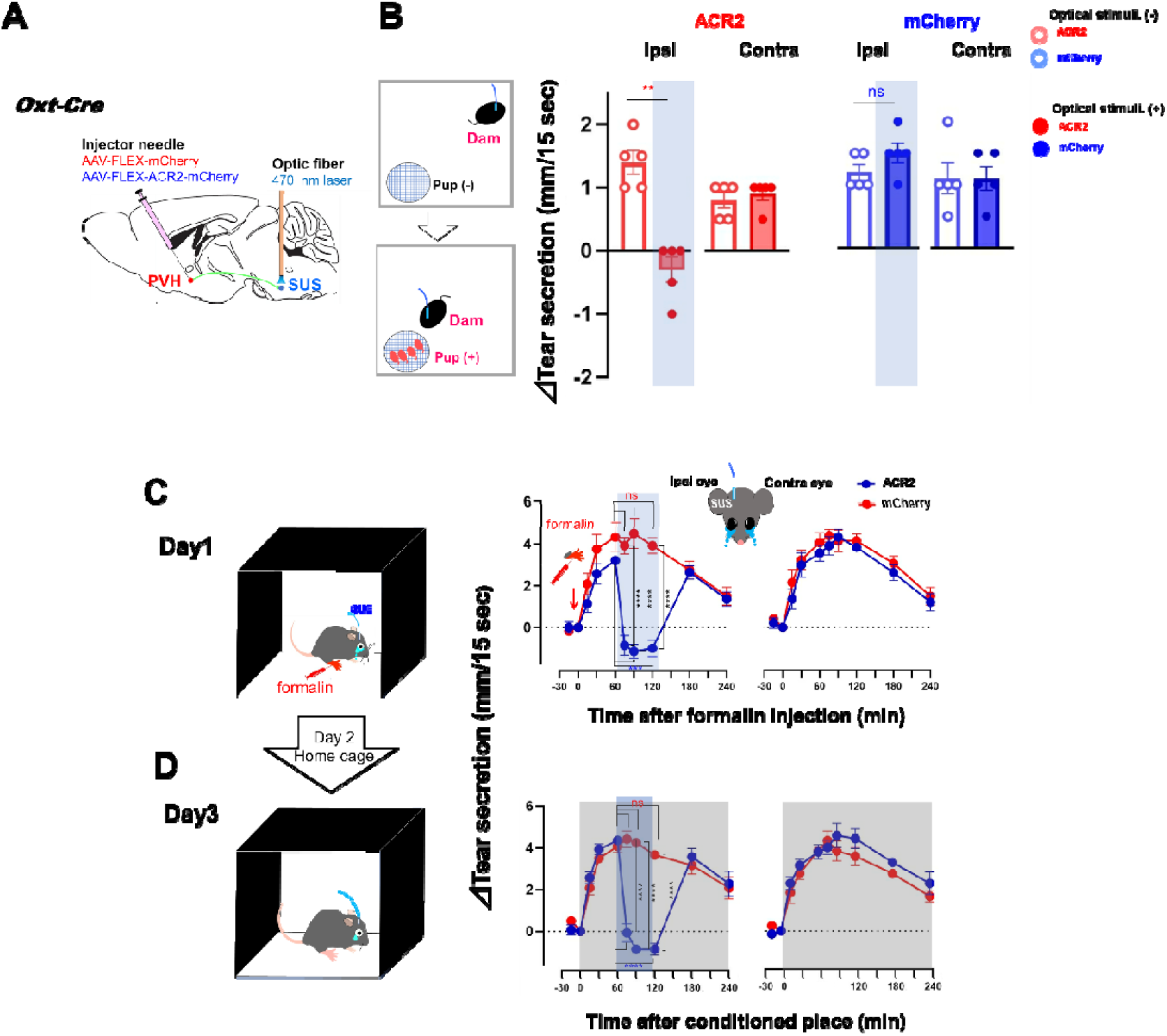
Mouse emotional behavior-linked tearing is suppressed by silencing of Oxt^PVH^→SUS neurons. A. Schematic of optogenetic inhibition of Oxt^PVH^→^SUS^ neurons during tearing. B. Changes in tear secretion (ACR2 n = 5, mCherry n = 5 per group) with optogenetic inhibition of Oxt^PVH^ → ^SUS^ neurons during mother pup reunion. The shaded area represents optical stimulation of the eye. C. Changes in tear secretion (mCherry n = 6, ACR2 n = 7per group) with optogenetic inhibition of Oxt^PVH^→^SUS^ neurons during tearing induced by PF (day 1). D. Changes in tear secretion (mCherry n = 6, ACR2 n = 7 per group) with optogenetic inhibition of Oxt^PVH^→^SUS^ neurons during re-exposure to the context condition (day 3). J. The shaded area represents the periods of optical stimulation (Fig C and D). \*\*\*\**P* < 0.0001, \*\**P* < 0.01.

It is unclear whether the functional involvement of Oxt^PVH^→^SUS^ in tearing as well as expression in OXTR^SUS^ is relevant to humans. Considering that the function of the central Oxt system lineage during evolution is also largely conserved across mammals (*36*), the central Oxt system reported in this study may have a common role in the regulation of tearing in humans, including emotionally evoked. The behavioral response to external physical (persistent nociception) or internal (Maternal and memory retrieval) emotional cues, which elicited tearing in mice in this study, is a fundamental reaction for survival across mammalian species. In addition, we have identified in a dog, a mammalian species reportedly having notable human-like social-cognitive skills, tearing evoked when reunited with the dog’s owner suggesting that non-human mammalian species, at least mouse and canine, have the potential for evoked emotional-linked tearing (*37*). Therefore, our study reveals that emotion-linked tearing is not a physiological reaction specific to humans (*Homo sapiens*) that was acquired during evolution but constitutes an evolutionarily conserved behavior originating from lower mammals and passed on to humans.

Despite substantial debate regarding the significance of emotional tears in humans from psychological and philosophical perspectives, the physiological role and interpretation of tearing in response to emotional arousal has been poorly investigated. In Charles Darwin’s book *Expression of the Emotions in Man and Animals* (1872) (*38*), emotional tearing is proposed to be an incidental result of reflex tearing, and this phenomenon is purposeless. Paradoxically, a recent study reported that tears contain chemosignals in both humans (*39*)(*40*)and mice (*41*)(*42*) that trigger social responses. Although we did not demonstrate direct evidence of the biological significance of emotional linked tearing, our findings, which demonstrate the involvement of the Oxt^PVH SUS^ and OXTR^SUS^ in ChAT ^SUS^ neuron in tear secretory neural pathway in emotion-linked tearing, facilitate a neuroscience-based approach that promotes elucidation of the meaning of emotional tears, thereby providing critical insight into the origin of emotion.

## Acknowledgements

We thank Dr Ayumu Inutsuka of Jichi Medical University for helpful advice regarding experiments. This experimental work was supported by a JST CREST grant (Grant Number JPMJCR1656 to A.Y) and JSPS KAKENHI grant (Grant Number18H02523 and 18KK0223 to A.Y., and 20K18392 and 17K16983 to T. I.).

## Author contributions

S.N., A.Y., K.F.T., T.K., and K.T. designed the research. A.Y. generated the virus vector. K.N. generated the OXTR-Venus mice. T.I. and Y.M. conducted calcium imaging experiments in acute brain slices. T.K., K.M., I.K., K.N., N.H., and N.K. conducted and analysed behaviour experiments. K.J., M.S., H.S., and F. I. assisted in the collection and analysis of data. S.N. and T.I. drafted the manuscript, and A.Y., K.F.T., K.M., T.K., M.M., and K.T. edited it. All authors read and approved the manuscript.

## Data and code availability statement

The data that support the findings of this study are available from the corresponding author on reasonable request.

## References

1. A. Jordan, J. Baum, Basic Tear Flow: Does It Exist? Ophthalmology. 87, 920–930 (1980).

2. A. Vingerhoets, Why Only Humans Weep: Unravelling the Mysteries of tears. Oxford Univ. Press. Oxford (2013) (available at http://ovidsp.ovid.com/ovidweb.cgi?T=JS&PAGE=reference&D=psyc10&NEWS=N&AN=2013-02266-000).

3. I. E. Toth, Z. Boldogkoi, I. Medveczky, M. Palkovits, “Lacrimal preganglionic neurons form a subdivision of the superior salivatory nucleus of rat: transneuronal labelling by pseudorabies virus” (1999).

4. S. Y. Botelho, TEARS AND THE LACRIMAL GLAND. Sci. Am. 211, 78–86 (1964).

5. J. C. Geerling, J. W. Shin, P. C. Chimenti, A. D. Loewy, Paraventricular hypothalamic nucleus: Axonal projections to the brainstem. J. Comp. Neurol. 518, 1460–1499 (2010).

6. R. Noseda, A. J. Lee, R. R. Nir, C. A. Bernstein, V. M. Kainz, S. M. Bertisch, C. Buettner, D. Borsook, R. Burstein, Neural mechanism for hypothalamic-mediated autonomic responses to light during Migraine. Proc. Natl. Acad. Sci. U. S. A. 114, E5683–E5692 (2017).

7. Y. Hosoya, M. Matsushita, Y. Sugiura, A direct hypothalamic projection to the superior salivatory nucleus neurons in the rat. A study using anterograde autoradiographic and retrograde HRP methods. Brain Res. 266, 329–333 (1983).

8. A. Argiolas, G. L. Gessa, Central Functions of Oxytocin. Neurosci. Biobehav. Rev. 15, 217–231 (1991).

9. S. Boll, A. C. Almeida de Minas, A. Raftogianni, S. C. Herpertz, V. Grinevich, Oxytocin and Pain Perception: From Animal Models to Human Research. Neuroscience. 387 (2018), pp. 149–161.

10. J. Bick, M. Dozier, K. Bernard, D. Grasso, R. Simons, Foster Mother-Infant Bonding: Associations Between Foster Mothers’ Oxytocin Production, Electrophysiological Brain Activity, Feelings of Commitment, and Caregiving Quality. Child Dev. 84, 826–840 (2013).

11. A. Argiolas, G. L. Gessa, “Central Functions of Oxytocin” (Pergamon Press plc, 1991).

12. P. J. Ryan, S. I. Ross, C. A. Campos, V. A. Derkach, R. D. Palmiter, Oxytocinreceptor-expressing neurons in the parabrachial nucleus regulate fluid intake. Nat. Neurosci. (2017), doi:10.1038/s41593-017-0014-z.

13. V. E. Chaves, C. Q. Tilelli, N. Almeida Brito, N. Brito, Role of oxytocin in energy metabolism. Peptides. 45, 9–14 (2013).

14. V. S. Fenelon, D. A. Poulain, D. T. Theodosis, Oxytocin neuron activation and fos expression: A quantitative immunocytochemical analysis of the effect of lactation, parturition, osmotic and cardiovascular stimulation. Neuroscience. 53, 77–89 (1993).

15. T. Kikusui, M. Kajita, N. Otsuka, T. Hattori, K. Kumazawa, A. Watarai, M. Nagasawa, A. Inutsuka, A. Yamanaka, N. Matsuo, H. E. Covington, K. Mogi, Sex differences in olfactory-induced neural activation of the amygdala. Behav. Brain Res. (2017), doi:10.1016/j.bbr.2017.11.034.

16. T. Kikusui, M. Kajita, N. Otsuka, T. Hattori, K. Kumazawa, A. Watarai, M. Nagasawa, A. Inutsuka, A. Yamanaka, N. Matsuo, H. E. Covington, K. Mogi, Sex differences in olfactory-induced neural activation of the amygdala. Behav. Brain Res. 346, 96–104 (2018).

17. K. K. Cover, B. N. Mathur, Axo-axonic synapses: Diversity in neural circuit function (2020), doi:10.1002/cne.25087.

18. S. Arrowsmith, S. Wray, Oxytocin: Its mechanism of action and receptor signalling in the myometrium. J. Neuroendocrinol. 26, 356–369 (2014).

19. D. Hawley, X. Tang, T. Zyrianova, M. Shah, S. Janga, A. Letourneau, M. Schicht, F. Paulsen, S. Hamm-Alvarez, H. P. Makarenkova, D. Zoukhri, Myoepithelial cell-driven acini contraction in response to oxytocin receptor stimulation is impaired in lacrimal glands of Sjögren’s syndrome animal models. Sci. Rep. 8, 9919 (2018).

20. K. Jin, T. Imada, R. Hisamura, M. Ito, H. Toriumi, K. F. Tanaka, S. Nakamura, K. Tsubota, Identification of Lacrimal Gland Postganglionic Innervation and Its Regulation of Tear Secretion. Am. J. Pathol. 190, 1068–1079 (2020).

21. H. S. Knobloch, A. Charlet, L. C. Hoffmann, M. Eliava, S. Khrulev, A. H. Cetin, P. Osten, M. K. Schwarz, P. H. Seeburg, R. Stoop, V. Grinevich, Evoked Axonal Oxytocin Release in the Central Amygdala Attenuates Fear Response. Neuron. 73, 553–566 (2012).

22. M. Yoshida, Y. Takayanagi, T. Onaka, K. Nishimori, Oxytocin receptor-Venus knock-in mice enable direct visualization of oxytocin receptor-expressing neurons. Neurosci. Res. 58, S222 (2007).

23. G. Gimpl, F. Fahrenholz, The oxytocin receptor system: Structure, function, and regulation. Physiol. Rev. 81, 629–683 (2001).

24. K. Horikawa, Y. Yamada, T. Matsuda, K. Kobayashi, M. Hashimoto, T. Matsu-ura, Miyawaki, T. Michikawa, K. Mikoshiba, T. Nagai, Spontaneous network activity visualized by ultrasensitive Ca2+ indicators, yellow Cameleon-Nano. Nat. Methods. 7, 729–732 (2010).

25. H. S. Knobloch, A. Charlet, L. C. Hoffmann, M. Eliava, S. Khrulev, A. H. Cetin, P. Osten, M. K. Schwarz, P. H. Seeburg, R. Stoop, V. Grinevich, Evoked Axonal Oxytocin Release in the Central Amygdala Attenuates Fear Response. Neuron. 73, 553–566 (2012).

26. P. J. Ryan, S. I. Ross, C. A. Campos, V. A. Derkach, R. D. Palmiter, Oxytocin-receptor-expressing neurons in the parabrachial nucleus regulate fluid intake. Nat. Neurosci. 20, 1722–1733 (2017).

27. J. R. Mutch, The Lacrimation reflex. Br. J. Ophthalmol. 28, 317–336 (1944).

28. M. E. Stern, J. Gao, K. F. Siemasko, R. W. Beuerman, S. C. Pflugfelder, The role of the lacrimal functional unit in the pathophysiology of dry eye. Exp. Eye Res. 78, 409–416 (2004).

29. K. Jin, T. Imada, S. Nakamura, Y. Izuta, E. Oonishi, M. Shibuya, H. Sakaguchi, H. Tanabe, M. Ito, K. Katanosaka, K. Tsubota, Corneal Sensory Experience via Transient Receptor Potential Vanilloid 1 Accelerates the Maturation of Neonatal Tearing. Am. J. Pathol. 189, 1699–1710 (2019).

30. J. Panksepp, The cross-mammalian neurophenomenology of primal emotional affects: From animal feelings to human therapeutics. J. Comp. Neurol. 524, 1624–1635 (2016).

31. J. LeDoux, Rethinking the Emotional Brain. Neuron. 73, 653–676 (2012).

32. D. Dubuisson, S. G. Dennis, The formalin test: a quantitative study of the analgesic effects of morphine, meperidine, and brain stem stimulation in rats and cats. Pain. 4, 161–174 (1977).

33. Y. Miyazawa, Y. Takahashi, A. M. Watabe, F. Kato, Predominant synaptic potentiation and activation in the right central amygdala are independent of bilateral parabrachial activation in the hemilateral trigeminal inflammatory pain model of rats, doi:10.1177/1744806918807102.

34. J. Bowlby, M. Fry, M. D. S. Ainsworth, W. H. Organization, Child care and the growth of love / by John Bowlbyl; abridged and ed. by Margery Fryl; with two new chapters by Mary D. Salter Ainsworth (1965) (available at https://apps.who.int/iris/handle/10665/39640).

35. A. Gračanin, L. M. Bylsma, A. J. J. M. Vingerhoets, Why Only Humans Shed Emotional Tears: Evolutionary and Cultural Perspectives. Hum. Nat. 29, 104–133 (2018).

36. Z. R. Donaldson, L. J. Young, Oxytocin, vasopressin, and the neurogenetics of sociality. Science (80-.). 322, 900–904 (2008).

37. K. Murata, M. Nagasawa, T. Onaka, N. Kanemaki, S. Nakamura, K. Tsubota, K. Mogi, T. Kikusui, bioRxiv, in press, doi:10.1101/2022.03.09.483532.

38. C. Darwin, The expression of the emotions in man and animals (Oxford University Press., 1872).

39. S. Gelstein, Y. Yeshurun, L. Rozenkrantz, S. Shushan, I. Frumin, Y. Roth, N. Sobel, Human tears contain a chemosignal. Science (80-.). 331, 226–230 (2011).

40. A. Gračanin, M. A. L. M. van Assen, V. Omrčen, I. Koraj, A. J. J. M. Vingerhoets, Chemosignalling effects of human tears revisited: Does exposure to female tears decrease males’ perception of female sexual attractiveness? Cogn. Emot. 31, 139–150 (2017).

41. S. Haga, T. Hattori, T. Sato, K. Sato, S. Matsuda, R. Kobayakawa, H. Sakano, Y. Yoshihara, T. Kikusui, K. Touhara, The male mouse pheromone ESP1 enhances female sexual receptive behaviour through a specific vomeronasal receptor. Nature. 466, 118–122 (2010).

42. M. Tsunoda, K. Miyamichi, R. Eguchi, Y. Sakuma, Y. Yoshihara, T. Kikusui, M. Kuwahara, K. Touhara, Identification of an Intra-and Inter-specific Tear Protein Signal in Rodents. Curr. Biol. (2018), doi:10.1016/j.cub.2018.02.060.

