## Supplementary material for "The oxytocin system regulates tearing": Supplemetal materials

**Supplemental materials**

Materials and Methods

Tables S1 to S25

Figures S1 to S16

Movies S1 to S4 legends

#### Materials and Methods

##### Animals

The following mouse lines were used in these experiments: heterozygous *Oxt-Cre* (024234, Jackson Laboratory), *OXTR-Cre* (030543, Jackson Laboratory), *OXTR-Venus* (MGI:3838764) (*1*), *ChAT-Cre* (006410, Jackson Laboratory), and wild type (C57BL/6J, Charles River Japan, Yokohama, Japan). *OxtCre/+::OXTRCre/+* mice were obtained by crossing *OxtCre/+* mice and *OXTRCre/+* mice (*2*). These transgenic mice were maintained on a C57BL/6J genetic background. We used both males and females in all experiments, 7–10 weeks old at start of experimentation with similar numbers where possible. Mice were housed in cages under the same conditions. Water and food were available ad libitum. No statistical methods were used to predetermine sample size.

All experimental protocols that involved animals were approved by the Animal Experimentation Ethics Committee of Keio University School of Medicine, Tokyo, Japan.

##### Tear secretion measurement

Clinically available cotton thread test was applied for the mice evaluation with some modifications (*3*). A phenol red thread (AYUMI Pharmaceutical Corporation, Tokyo, Japan) was placed on the temporal side of the upper eyelid margin for 15 seconds. The length of the moistened area from the edge was measured with a precision of 0.5 mm. Mice were allowed to be handled gently for acclimation at least 1 week before measurements.

##### Adeno-associated viral (AAV) vectors

We used recombinant AAV vectors with the FLEX switch system to specifically express ChR2, ACR2, hM3Dq, hM4Di, YCnano50 (Calcium imaging) or Flippase-Tetanus toxin C-fragment (Flp-TTC) in Cre-expressing neurons or the dFRT cassette for Flp-dependent hM3Dq expression control. Characteristics of AAVs are summarized in Table S1.

Table S1 Characteristics of AAV vectors

| AAV Plasmids | Serotype | Promoter and Expression Characteristics | Fluorescent labeled | Cre or Flp dependent | Activities | Titer (genome copies/ml) |
|---|---|---|---|---|---|---|
| AAV-CAG-FLEX-mCherry | AAV-9 | CAG (general) | mCherry | Cre | - | $2\times10^{12}$ |
| AAV-CMV-FLEX-ACR2-2A-mCherry | AAV-DJ | CMV (general) | mCherry | Cre | Optogenetic inhibition | $2\times10^{12}$ |

| | | | | | | |
|---|---|---|---|---|---|---|
| AAV-CMV-FLEX-ChR2(ET/TC)-EYFP | AAV-9 | CMV (general) | EGFP | Cre | Optogenetic activation | $6\times10^{11}$ |
| AAV-CMV-FLEX-ChR2(ET/TC)-mKate2 | AAV-9 | CMV (general) | mKate2 | Cre | Optogenetic activation | $3\times10^{12}$ |
| AAV-CAG-FLEX-mCherry-2A-Flp-TTC | AAV-DJ | CAG (general) | mCherry | Cre | Transsynaptic transportation of Flp | $3\times10^{12}$ |
| AAVCMV-dFRT-hM3Dq-mCherry-WPRE | AAV-DJ | CMV (general) | mCherry | Flp | rM3D(Gq) - activation | $3\times10^{12}$ |
| AAVCMV-FLEX-YCnano50 | AAV-DJ | CMV (general) | - | Cre | Yellow Cameleon nano50 expression | $1\times10^{13}$ |

**Stereotaxic surgeries**

Stereotaxic surgeries were performed at PVH and/or SUS under anesthesia using a combination of medetomidine (0.75 mg/kg), butorphanol (5 mg/kg), and midazolam (4 mg/kg).

The AAVs (600 nL) were infused with a glass micropipette (Harvard Apparatus, Holliston, MA, USA) using an air pressure injector (BEX, MODEL BJ110, Tokyo, Japan). Glass micropipettes were removed 5 min after infusions were complete. Optic fibres (ferrules) for the optogenetic manipulations and/or infusion cannulas (24-gauge guide cannulas sealed with dummy cannula) for the chemogenetic or pharmacologic stimulation were implanted above the selected coordinates. Fibres and cannulas were implanted and attached to the skull by dental cement (Sun Medical, Shiga, Japan) one week before the optogenetic manipulation, chemogenetic stimulation, or pharmacologic stimulation. Injection of AAV and implantation of fibres and cannulas were performed according to the coordinates in Table 2 based on standard stereotaxic coordinates (*4*).

Table S2 Stereotaxic Coordinates

| | Coordinates | AAV infusion | Optic fibres | Infusion |
|---|---|---|---|---|

| | | | | |
|---|---|---|---|---|
| | | | | cannulas |
| PVH | Anteroposterior (AP) | -0.8 mm | | |
| | Mediolateral (ML) | ±0.2 mm | | |
| | Dorsoventral (DV) | -4.7 mm | -4.6 mm | -4.5 mm |
| SUS | AP | -5.6 mm | | |
| | ML | ± 1.5 mm | | |
| | DV | -5.3 mm | -5.2 mm | -5.1 mm |

**Neuronal manipulations**

**Optogenetic manipulation**

A ceramic optic fibre ferrules (core-diameter, 0.22 NA; KYOCERA, Kyoto, Japan) were implanted above ipsilateral PVH or SUS. Optical stimulation was performed with a fibre-coupled LED with peak emission at 470 nm (M470F3, ThorLabs, Newton, NJ, USA) for 10 to 20 minutes. LED intensity was controlled using a LED driver with analogue modulation (LEDD1B, ThorLabs) connected to a pulse generator (FG085 mini DDS function generator, JYE Tech, Guangxi, China). The maximal LED light power intensity at the tip of the optical fibre was 0.8 mW. Stimulation power was varied between 0.8 mW (high) and 0.2 mW (low) of light power for the inhibition and 5 Hz (low) and 10 Hz (high) of light frequency for the activation, respectively.

**Chemogenetic manipulation**

For the activation of hM3Dq-expressed $Oxt^{PVH \rightarrow SUS}$ neurons, CNO was intraperitoneally injected or intra-PVH infused through infusion cannula at the dose of 180 ng or 1 mg/kg, respectively. CNO was infused at a rate of 1 μL per minutes using microinjection pump (YMC, MODEL YSP-101, Kyoto, Japan) with Hamilton syringe (Hamilton, 1002TLL 2.5 ml SYR, Reno, NV, USA) for 1 minute. CNO was dissolved to the desired concentration with saline and aCSF and used as control for intraperitoneal injection and intra-PVH infusion, respectively.

**Pharmacologic manipulation**

SUS was blocked by OXTR antagonist L-368,899 (0.004-40 ng, Tocris Bioscience, Bristol, UK) and gutamate channel blockers, amino-3-hydroxy-5-methylisoxazole-4-propionic acid (AMPA) receptor antagonist AP-5 (0.01 ng, Cayman Chemical, Ann Arbor, MI, USA) or N-methyl-D-aspartate receptors antagonist NBQX disodium salt (0.05 ng, Abcam, Cambridge, MA, USA) under the activation of $Oxt^{PVH}$ neurons. Antagonists were infused one minute before optical stimulation of $Oxt^{PVH}$.

SUS was activated by Oxt (0-400 ng) or OXTR agonist Way 267,464 (0-400 ng, Tocris

Bioscience, Ellisville, MO, USA) agonist, and the antagonist was diluted with aCSF and used as a control. Each solution was infused into SUS through an infusion cannula under the same condition of intra-PVH infusion.

#### Lacrimal nerve denervation

Surgical denervation of the lacrimal nerve innervated in the LG was performed according to our previous report (*3*). In brief, mice were placed in a prone position, and skin on the ipsilateral temporal side of the head was incised under deep anaesthesia using a combination anaesthetic. The post-ganglionic nerve bundle was detached from the blood vessels at the caudal root site of the ventral surface of the LG and denervated under a stereomicroscope. Denervation was performed only on the ipsilateral LG, and the contralateral LG remained as a control. Lacrimal nerve was denervated before one day of optogenetic stimulation of $Oxt^{PVH}$.

#### *In vivo* LG perfusion

The mice were first anaesthetised with an intraperitoneal injection of a combination anaesthetic. They were then placed in a lateral position, and their skin on the left temporal side of the head was incised. After the LG was exteriorised, a custom-built ring was attached to the masseter muscle using a cyanoacrylate-based glue (Loctite, Henkel Japan, Yokohama, Japan) surrounding the LG. The perfusion chamber was fitted on the ring fixed to a head-holder. The LG was continuously perfused with saline solution (in mM: 140 mM NaCl, 5 mM KCl, 2 mM $CaCl_2$, 1 mM $MgCl_2$, 10 mM HEPES, and 10 mM dextrose; pH 7.4) at a flow rate of 0.8 mL/minute. For the conformation of LG activity, Oxt (20 nM) or Acetyl choline chloride (1 μM ) For the optical stimulation of $Oxt^{PVH}$, OXTR antagonist atosiban (100 nM) perfusion was started 30 sec before stimulation and continued for 10 minutes.

#### Histology

##### Brain sections preparation

Mice were anaesthetised with a combination anaesthetic, and perfused transcardially with chilled 4% paraformaldehyde (PFA). The brains were removed and were post-fixed in 4% PFA at 4°C overnight and then soaked in 30% sucrose for 48 hours. Coronal sections were cut with a cryostat and immersed in blocking buffer (1% bovine serum albumin [BSA] and 0.25% Triton-X in PBS), then incubated with primary antibodies at 4°C, overnight. The sections were washed with blocking buffer, then incubated with secondary antibodies for 1 hour at room temperature. The brain sections were mounted and stored at 4°C, until observation. Brain sections were immuno-stained with free-floating (PVH, 60 μm) or slide-mounted (SUS, 25 μm) manner.

**Antibodies**

The $Oxt^{PVH}$ neurons were stained with mouse anti-neurophysin I antibody (1:300,Sigma-Aldrich, clone PS38) (*5*). To detect $OXTR^{SUS}$ neurons, fluorescent protein Venus or GFP expressed under the control of OXTR promotor in *OXTR Venus* mouse and *OXTR-Cre* mice were stained with rabbit anti-GFP (1:300, MBL, Woburn, MA, USA) and mouse anti-GFP (1:300, Fujifilm Wako Pure Chemical, Osaka, Japan), respectively. The SUS ChAT neurons were stained with goat anti-ChAT (1:100, Millipore, Watford, UK). To amplify the signal of Cre-dependent fluorescent protein expressed by AAV vectors, mouse anti-GFP, rabbit anti-GFP, rabbit anti-tRFP (1:400, Evrogen JSC, Moscow, Russia), goat anti-mCherry (1:500, Origene Technologies, Rockville, MD, USA) and rabbit anti-mCherry (1:500, Abcam) were used. For the c-Fos study, mice were perfused after 1.5 hours of optical stimulation (10 Hz, 10 min) or hind paw injection of formalin or saline, and their brain sections were stained with guinea pig anti-cFOS (1:300, Synaptic Systems, Goettingen, Germany). Secondary antibodies were optimally selected from the following; Alexa Fluor 488-conjugated donkey anti-mouse, anti-rabbit, anti-goat, and anti-guinea pig antibody (1:300, Invitrogen; Jackson Immunoresearch Laboratories, West Grove, PA, USA); Alexa Fluor 555-conjugated donkey anti-mouse and anti-goat antibody (1:300, Invitrogen); and Alexa Fluor 647-conjugated donkey anti-rabbit and anti-goat antibody (1:300, Invitrogen).

**Microscopic observation**

Brain sections were observed and imaged with a confocal microscopy (LSM710: Carl Zeiss, Germany, FV3000: Olympus, Tokyo, Japan) or a fluorescence microscope (BZ9000, Keyence, Osaka, Japan). Images were minimally processed to enhance brightness and contrast for optimal presentation using ZEN (Carl Zeiss Microscopy), FV31S (Olympus), or BZ-Analyzer. (Keyence).

**Cell counting**

For cell counting, a series consisting of two sections of the SUS and four sections of the PVH were examined for each subject.

**Acute brain slice $Ca^{2+}$ imaging of SUS**

**Acute brain slice**

$Ca^{2+}$ indicator YCnano50 was specifically expressed in the SUS ChAT neurons by injecting AAV- FLEX-YCnano50 into the SUS of *ChAT-Cre* mice. After 3 weeks of AAV injection, acute 300 μm coronal brain section including SUS were prepared in ice-cold cutting solution (in mM: 110 K-gluconate, 15 KCl, 0.05 EGTA, 5 HEPES, 26.2 $NaHCO_3$,

25 Glucose, 3.3 $MgCl_2$ and 0.0015 (±)-3-(2-Carboxypiperazin-4-y1)-1-phosphonic acid) with a rotorslicer DTY-8700 (Dosaka EM, Osaka, Japan). The slices were temporarily placed in a bath solution (aCSF, in mM: 124 NaCl, 3 KCl, 2 $MgCl_2$, 2 $CaCl_2$, 1.23 $NaH_2PO_4$, 26 $NaHCO_3$ and 25 Glucose) at 35°C for 1 hour and then incubated in the same solution at room temperature for another 1 hour. The slices were constantly gassed with 95% $O_2$ and 5% $CO_2$.

##### Two photon $Ca^{2+}$ imaging

Brain slices were transformed to a recording chamber (RC-26; Warner Instruments, Hamden, CT, USA) on a two photon microscope (FV1200MPE, Olympus, Tokyo, Japan) equipped with a water immersion objective lens (XLPlaN25 × 1.05WMP, Olympus) which connected to a femtosecond laser source, Ti:sapphire laser (MaiTaiHP: Spectra Physics, Santa Clare, Ca, USA). Slices were constantly perfused with bath solution gassed with 95% $O_2$ and 5% $CO_2$. Neurons of interest were identified by CFP and YFP fluorescence of YCnano50. For observation of the $Ca^{2+}$ signal, an excitation wavelength of 870 nm was used, and two-photon excited fluorescence images of CFP and YFP were acquired in separate channels through dichroic mirrors and emission filters, BP460-500 and BP520-560, respectively. Images were acquired at 2 sec. per frame, and the obtained images were analysed with the Aqua Cosmos software (Hamamatsu Photonics, Shizuoka, Japan).

##### Pharmacological treatments

Solutions of Oxt (0.05, 0.5, and 5 µM; Peptide Institute, Osaka, Japan), OXTR agonist Way-267,464 (5 µM, Tocris Bioscience), OXTR antagonist L-368,899 (5 µM), and high-$K^+$ (50 mM potassium chloride) were diluted to the desired concentrations with the 1 µM TTX-containing bath solution. Oxt and Way-267,464 were applied to the brain slice for 2 minutes. To block the OXTR, L-368,899 was applied to brain slices 5 minutes before the application of 5 µM Oxt or Way-267,464 and lasted for 9 minutes.

#### Induction of reflex tearing

Mice were treated with eye drops of 1 µM capsaicin (capsaicin diluted with 0.095% ethanol in physiological saline, Wako Pure Chemical, Osaka, Japan). During optical inhibition of $Oxt^{PVH \rightarrow SUS}$ or $OXTR^{SUS}$, a 1 µL drop of capsaicin or vehicle solution was softly dropped to the cornea of bilateral eye using a micropipette. Immediately after eye drops instillation, excess solutions in eyelid margin was gently wiped off with a swab.

#### Behavioral procedures

Mice were exposed to the following positive or negative valence emotional cues. Changes in tear secretion were measured before and up to 60 min after exposure and compared with those of non-treated mice.

**Isolation and reunion with cage mates**

Two mice were housed together in a conventional plastic cage for at least 3−5 weeks. Then, one mouse was isolated in the same plastic cage for 1 hour in another room (alone). After 1 hour of isolation, mice were returned to their home cages with residents (reunion).

**Tail suspension**

Mice that were both acoustically and visually isolated were suspended 50 cm above the floor with adhesive tape placed approximately 1 cm from the tip of the tail for 6 min (*6*).

**Encounter with an intruder**

Mice were conditioned in a plastic cage for at least 2 weeks, and CD1 mice were placed into the home cage of residents for 10 min.

**Suckling mother**

Suckling dams within 6 days postpartum were used for tear secretion measurements.

**Enriched environmental rehousing**

Mice were individually conditioned in a plastic cage with a voluntary running wheel and shelter for 1 week. The mice were then housed without enrichment devices for 1 day. Changes in tear secretion were measured after the enrichment devices were replaced in their cages.

**Drinking water re-supply or preferred food re-feeding elicited relief**

Mice were water-deprived 24 h before the experiment. Changes in tear secretion were measured 30 min after *ad libitum* access to water was re-established (WR). Lab chow was made available along with chocolate (2 ± 0.5 g) (milk chocolate, Meiji) for 1 week. After 1-day deprivation of chocolate, changes in tear secretion were measured after chocolate re-feeding(*7*).

**Electric foot shock**

Mice were subjected to a single electric foot shock (0.40 mA for 2 s) using the PRECISION ANIMAL SHOCKER 110 V (Coulbourn Instruments, Allentown, PA, USA; E13-14; EFS).

**Hot plate placing**

Mice were placed on a hot-plate apparatus, which was maintained at 50 ± 0.1°C. Mice were returned to their home cages immediately after nociceptive behaviors, such as hind-paw lifting, licking, and jumping, were observed.

**Tail pinching**

Single pinch stimulation with forceps was applied to the tails of mice at a force of 300 *g*.

**Facial formalin injections**

Mice were injected with 5 µL of 5% formalin (37% formaldehyde solution (Nacalai Tesque Inc.) diluted with saline (0.9% NaCl) into the upper lip with a microsyringe with a 30-gauge needle attached.

**Paw formalin injection**

Mice were injected with 5 µL of 5% formalin into the hind paw using a microsyringe with a 30-gauge needle attached.

**Nociceptive behavior evaluation**

The sum of time spent exhibiting characteristic nociceptive behaviors, such as paw lifting, licking, and biting of the formalin-injected paw was calculated 120 min after injection.

**Mother pup separation and reunion**

Postpartum dams within 6 days of suckling more than four pups were used. Dams were separated from their pups for 2 h before reunion by removing the pups from the home cage and placing them in a fresh cage. Pups were housed in a wire mesh ball (5 cm diameter) placed in the home cage during the separation period and were reunited with dams to restrict direct interactions. Tear secretions were measured during separation and reunion periods. Dams' pup retrieval behavior, frequency of nose contact with the pup area, and distance moved were analyzed during the reunion period using EthoVision. XT (Noldus, Wageningen, The Netherlands). For virgin females, other pups were encountered in the same manner as that for suckling dams.

**Paw formalin injections**

Mice were injected with 5 µL of 5% formalin into the hind paw using a microsyringe with a 30-gauge needle attached. To examine the relationship between tearing and nociceptive responses, mice were injected with 5 µL of 0, 2.5, and 5% formalin, and tear secretion was measured for up to 240 min.

**Aversive memory conditioning and retrieval**

For aversive memory conditioning, paw formalin injections were performed in a context conditioning chamber (black and white Plexiglas walls: 17 × 17 × 27 cm high). Mice were housed in the chamber for 240 min. Retrieval and extinction of aversive memories was performed by re-exposing mice to the same context chamber for 240 min without formalin injections at 3, 6, and 8 days after conditioning. Tear secretion was measured for 240 min. Immobilization time was analyzed using Any-maze (Stoelting Co, IL, USA) video tracking software.

To assess whether changes in tear secretion were associated with the context condition, re-exposure to the contextual condition was compared between three groups: (i) the home cage chamber with bedding (HC), two-chambered preference apparatus in which the context chamber was connected with (ii) the same chamber (CC-CC), and (iii) the home cage chamber with bedding (CC-HC) separated by a removable guillotine door to confine the animal. The guillotine separation door was opened 30 min after re-exposure to allow the mouse to passively cross the apparatus.

**Statistical analyses and figure preparation**

Data were first assessed for whether each data set passed the Shapiro-Wilk normality test and that the variance for each variable was similar across groups. Then, appropriate parametric or non-parametric statistical tests were used to make comparisons between groups. All analyses and plots were conducted using Prism (version 8.42 for windows, GraphPad Software, San Diego, CA, USA). Results are presented as standard error of the mean (SEM). Further details about the statistical tests are provided in the supplemental tables.

**Figure S1**

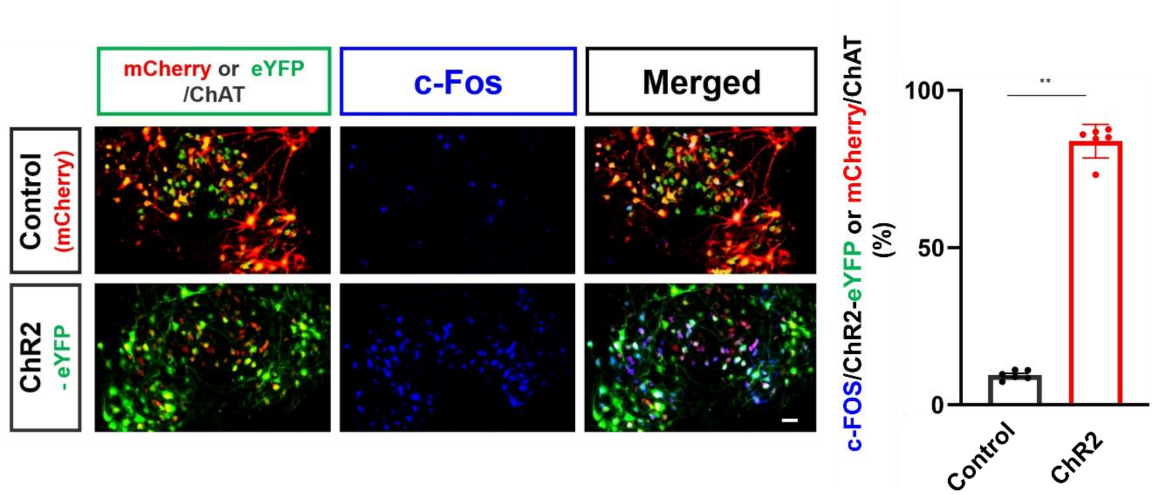


**Figure S1. Superior salivatory nucleus c-Fos expression ratio in mCherry- or ChR2-expressing ChAT$^{SUS}$ neurons after optical stimulation.**

The right bar chart shows the percentage of the mCherry- or ChR2- eYFP expressing ChAT$^{SUS}$ neurons merged with c-Fos. n = 6 per group. Scale bar represents 50 μm. ****$P < 0.0001$. See Table S13 for the statistical analyses.

**Figure S2**

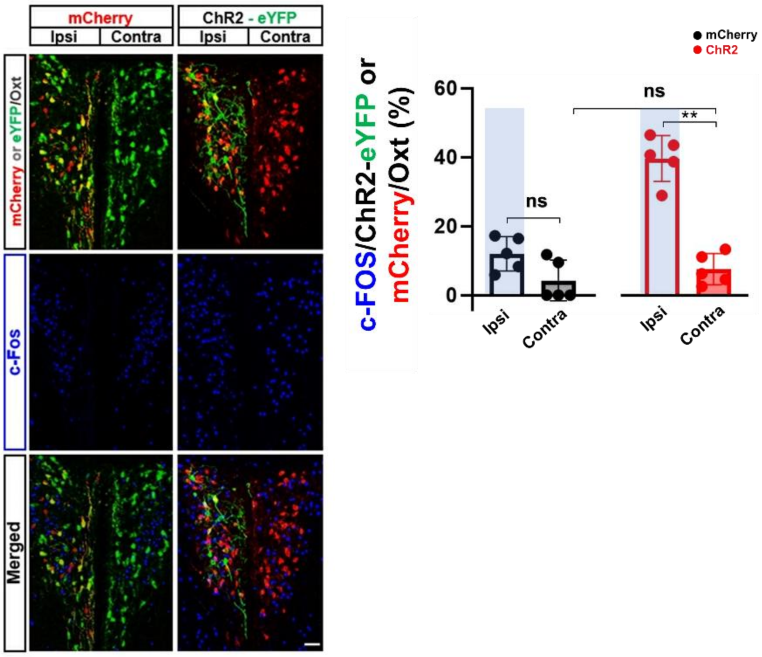


**Figure S2. Paraventricular hypothalamus c-Fos expression ratio in mCherry- or ChR2-expressing Oxt$^{PVH}$ neurons after optical stimulation.**

The right bar chart shows the percentage of the mCherry- or ChR2- eYFP expressing Oxt$^{PVH}$ neurons merged with c-Fos. Significantly large c-Fos expression were observed in the ipsilateral ChR2-eYFP expressed Oxt$^{PVH}$ compared to that of the contralateral and ipsilateral mCherry expressed Oxt$^{PVH}$.

n = 5 per group. Scale bar, 100 μm. The shaded areas in the figure represent the side of optical stimulation. ****$P < 0.0001$. See Table S13 for the statistical analyses.

**Figure S3**

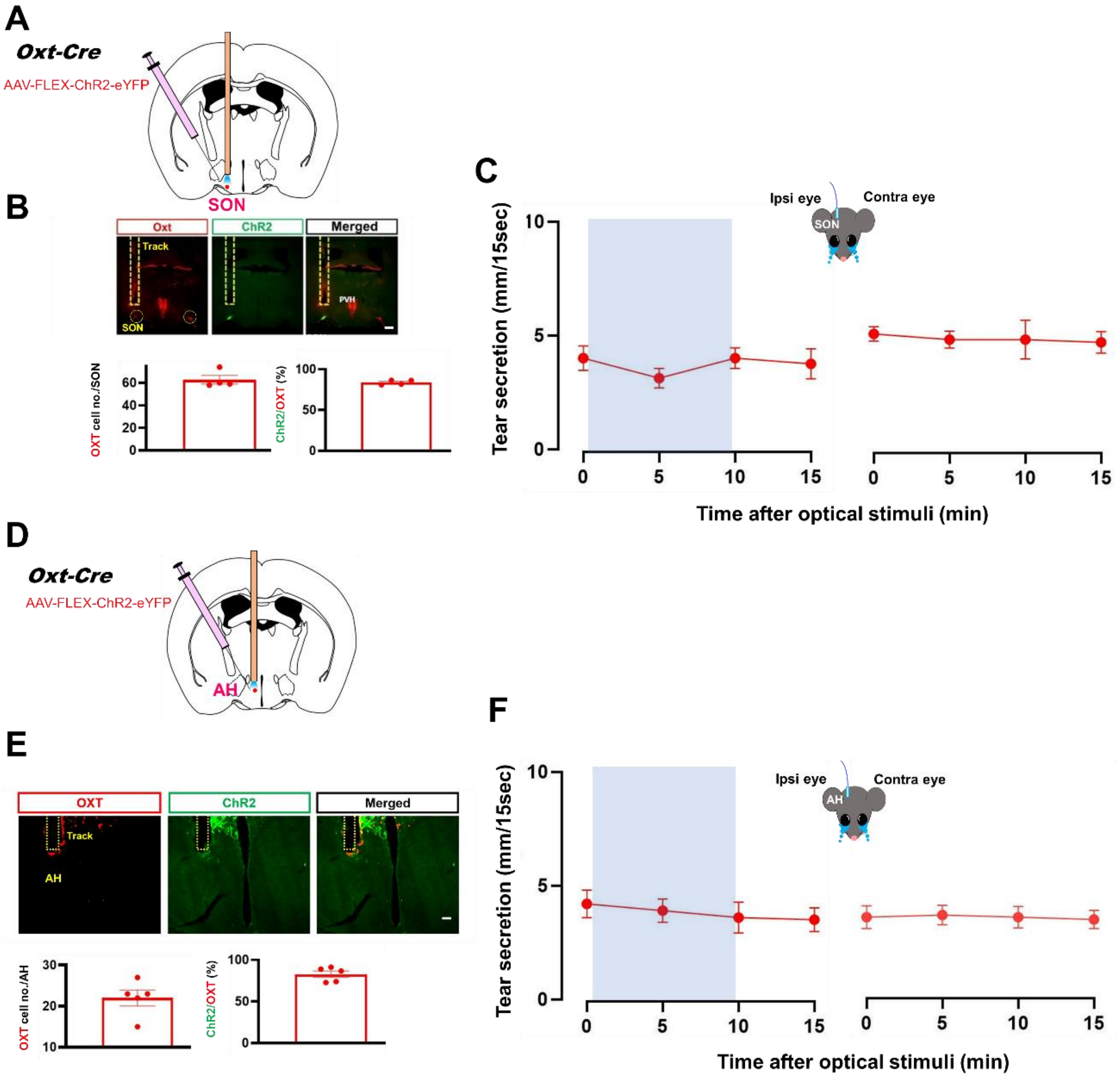


**Figure S3. Optical stimulation of SON and AH OXT neurons did not alter tear secretion.**

A. Schematic of optogenetic activation of $OXT^{SON}$ neurons.

B. Histochemical confirmation of ChR2 in $OXT^{SON}$ neurons.

C. Changes in tear secretion (n = 4 per group).

D. Schematic of optogenetic activation of $OXT^{AH}$ neurons.

E. Histochemical confirmation of ChR2 in $OXT^{AH}$ neurons.

F. Changes in tear secretion (n = 5 per group).

n = 4 (B, C) and 5 (E, F) per group. Scale bar represents 100 μm.

The shaded areas in the figures represent the periods (C and F) of optical stimulation. See Table S14 for the statistical analyses.

### Figure S4

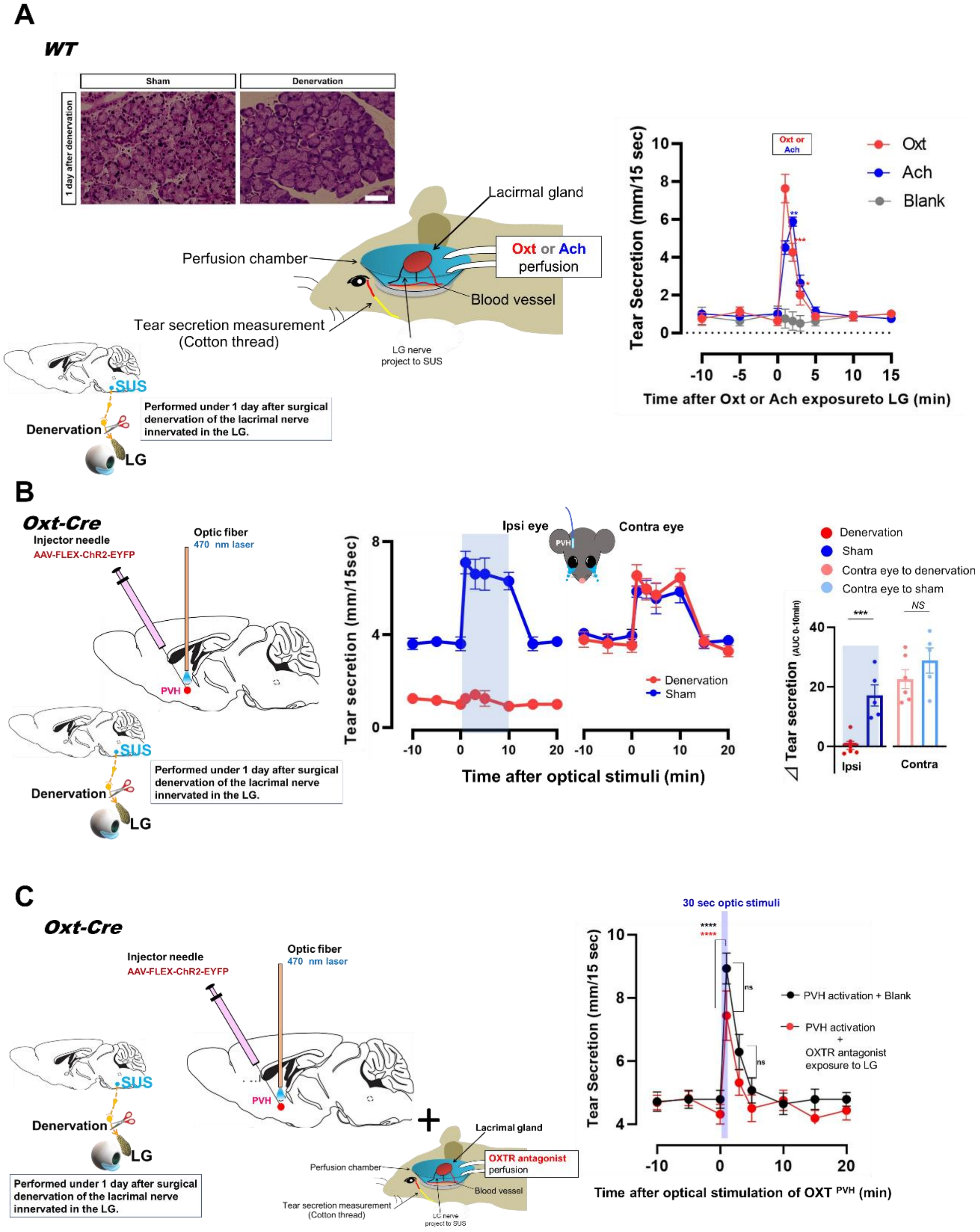

A
WT
Sham
Denervation
1 day after denervation
Lacirmal gland
Perfusion chamber
Oxt or Ach perfusion
Blood vessel
Tear secretion measurement (Cotton thread)
LG nerve project to SUS
SUS
Denervation
LG
Performed under 1 day after surgical denervation of the lacrimal nerve innervated in the LG.
Tear Secretion (mm/15 sec)
Oxt or Ach
Oxt
Ach
Blank
Time after Oxt or Ach exposureto LG (min)
B
Oxt-Cre
Injector needle
AAV-FLEX-ChR2-EYFP
Optic fiber
470 nm laser
PVH
SUS
Denervation
LG
Performed under 1 day after surgical denervation of the lacrimal nerve innervated in the LG.
Tear secretion (mm/15sec)
Ipsi eye
Contra eye
Denervation
Sham
Time after optical stimuli (min)
Denervation
Sham
Contra eye to denervation
Contra eye to sham
***
NS
⊿ Tear secretion (AUC 0-10min)
Ipsi
Contra
C
Oxt-Cre
Injector needle
AAV-FLEX-ChR2-EYFP
Optic fiber
470 nm laser
PVH
SUS
Denervation
LG
Performed under 1 day after surgical denervation of the lacrimal nerve innervated in the LG.
Lacrimal gland
Perfusion chamber
OXTR antagonist perfusion
Blood vessel
Tear secretion measurement (Cotton thread)
LG nerve project to SUS
30 sec optic stimuli
Tear Secretion (mm/15 sec)
PVH activation + Blank
PVH activation + OXTR antagonist exposure to LG
ns
Time after optical stimulation of OXT PVH (min)

**Figure S4. Changes in the tear secretion by activation of the Oxt$^{PVH}$ neurons upon surgical denervation of the lacrimal nerve innervating the LG.**

A. Confirmation of the morphological alterations of LG acinar cells (upper left) or increase in the tear secretion by the application of Oxt or acetylcholine to the LG upon surgical denervation of the lacrimal nerve innervated in the LG (right, n = 4 per group). Confirmations were established 1 day after surgical denervation. Scale bar represents 50 μm.

B. Dynamics (left) and $\Delta AUC_{(0\text{-}10\ min).}$ (right) of the effect of lacrimal nerve denervation on the increase in tear secretion with optogenetic activation of Oxt$^{PVH}$ neurons (n = 5 per group). The shaded areas in the figures represent the periods (center line graph) or side (right bar chart) of optical stimulation.

C. Effect of exposure of OXTR antagonist to Oxt-Cre lacrimal gland during optical activation of Oxt$^{PVH}$ neurons (n = 7 per group). The shaded areas in the line figure represent the periods of optical stimulation. See Table S15 for the statistical analyses.

**Figure S5**

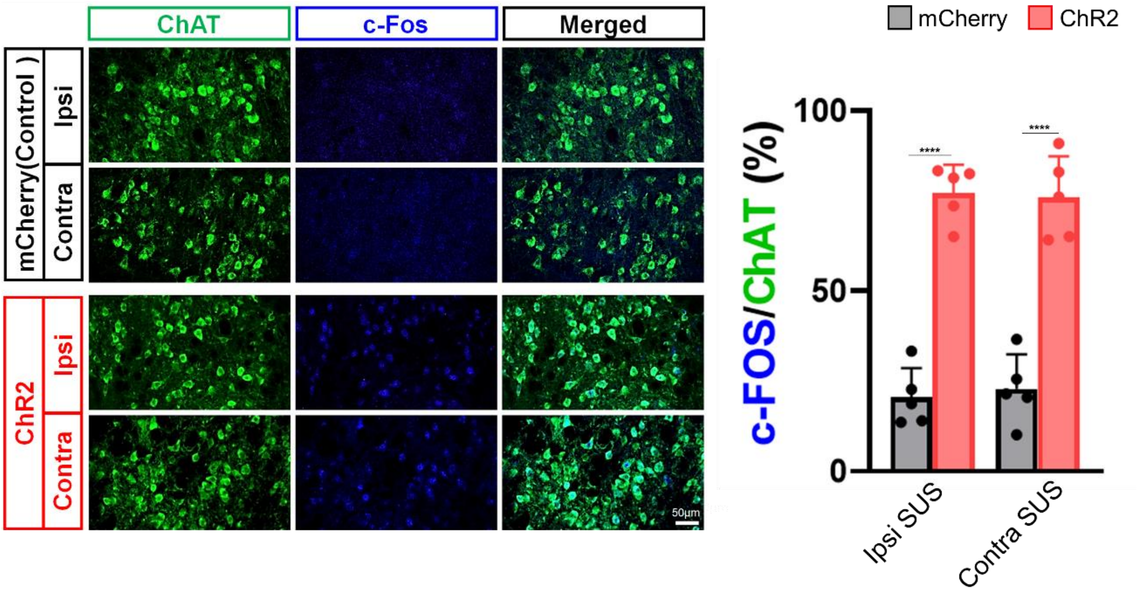


**Figure S5. Change in the SUS c-Fos expression by activation of Oxt$^{PVH}$ neurons.**
Scale bar represents 50 μm. The right bar chart shows the percentage of the ChAT$^{SUS}$ cells merged with c-Fos. n = 5 per group.
*****P* < 0.001. See Table S16 for the statistical analyses.

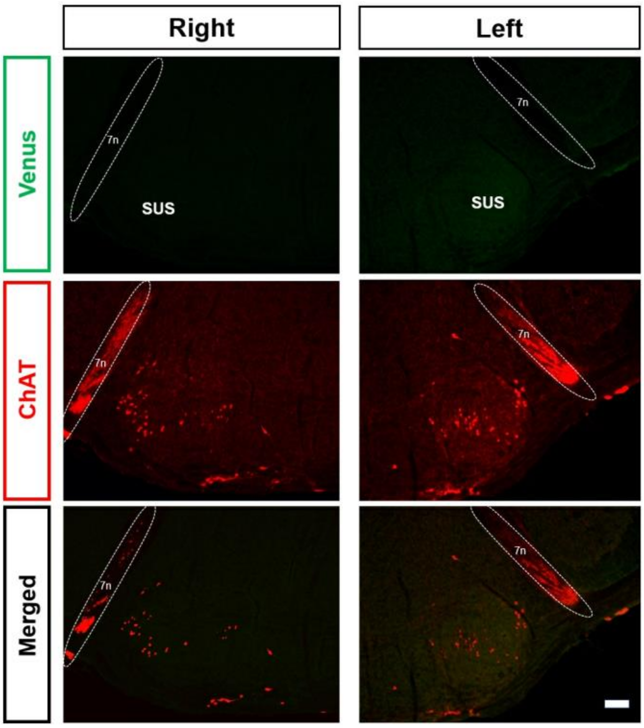


**Figure S6. Venus fluorescent protein expression in the SUS of OXTR$^{-Venus\ (-/-)}$ mice.**
Scale bar represents 100 μm.

**Figure S7**

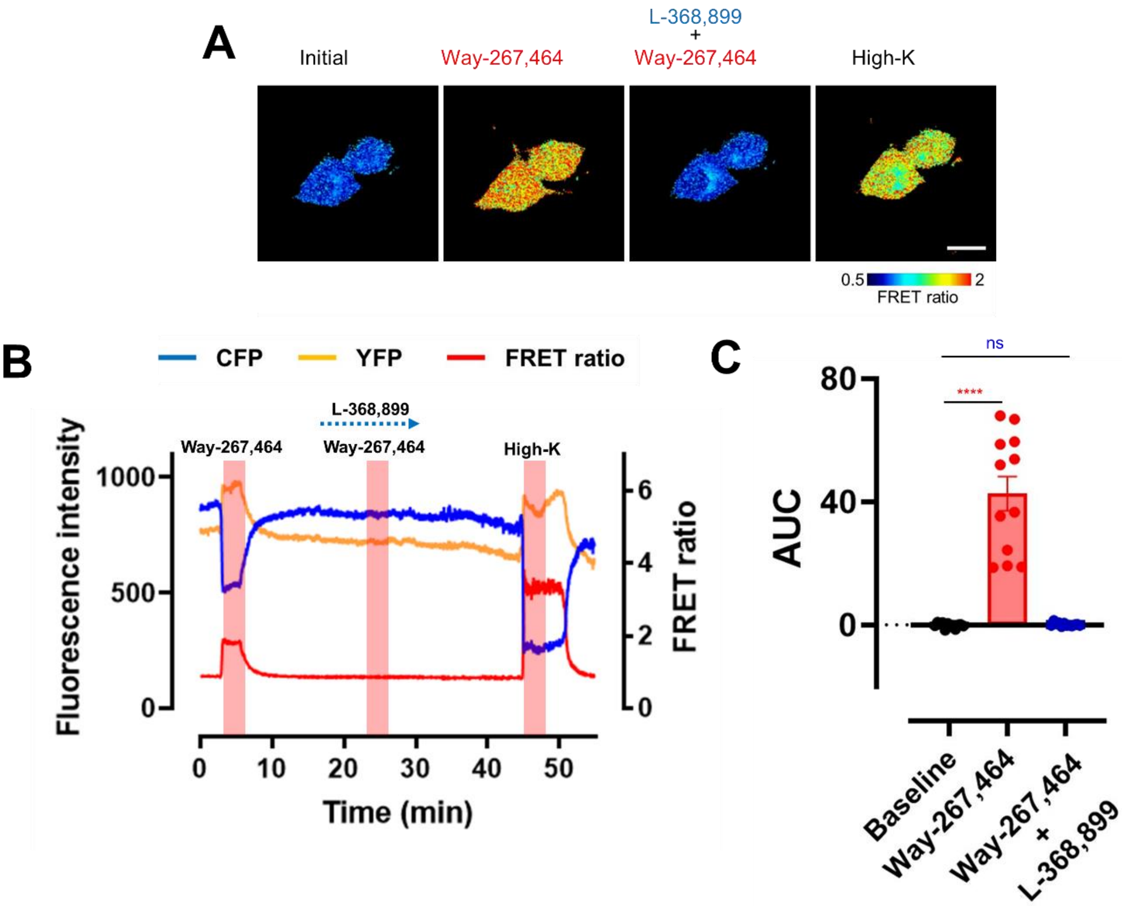


**Figure S7. Changes in the FRET ratio dynamics in SUS ChAT neurons stimulated by OXTR agonist Way-267,464 with/without OXTR antagonist L-368,899.**

A. Pseudo-color images of FRET ratio elevation in SUS ChAT neurons stimulated by OXTR agonist Way-267,464 with/without OXTR antagonist L-368,899 and high $K^+$. Scale bar represents 20 µm.

B. Representative fluorescence intensity of cyan fluorescent protein (CFP), YFP, and FRET ratio.

C. The AUC of the changes in the FRET ratio stimulated by OXTR agonist Way-267,464 with/without OXTR antagonist L-368,899. n = 12 neurons.

****$P < 0.001$. See Table S17 for the statistical analyses.

**Figure S8**

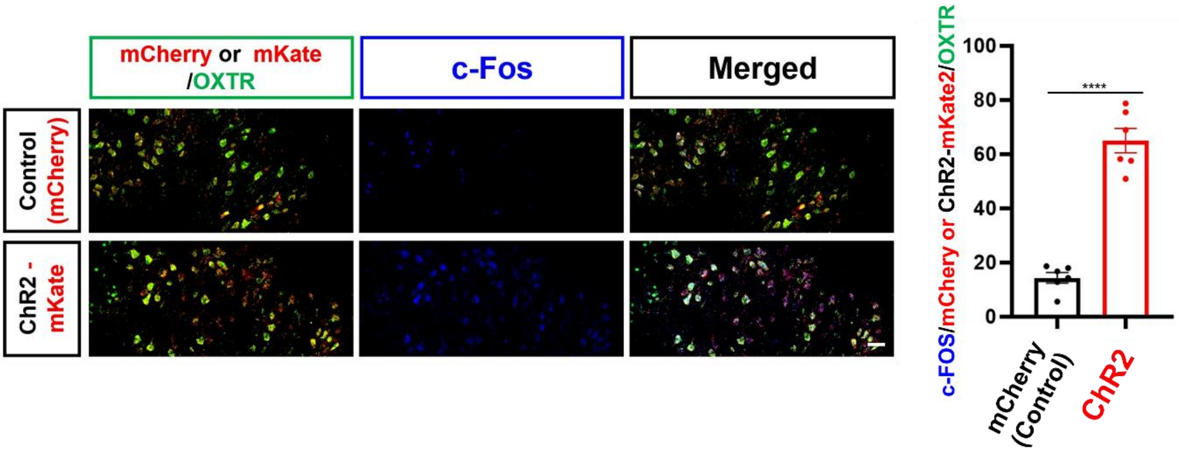


**Figure S8. Superior salivatory nucleus c-Fos expression ratio in mCherry- or ChR2-expressing OXTR$^{SUS}$ neurons after optical stimulation.**

The right bar chart shows the percentage of the mCherry- or ChR2- mKate2 expressing the OXTR$^{SUS}$ neurons merged with c-Fos. n = 6 per group. Scale bar represents 50 μm. ****$P$ < 0.001. See Table S18 for the statistical analyses.

**Figure S9**

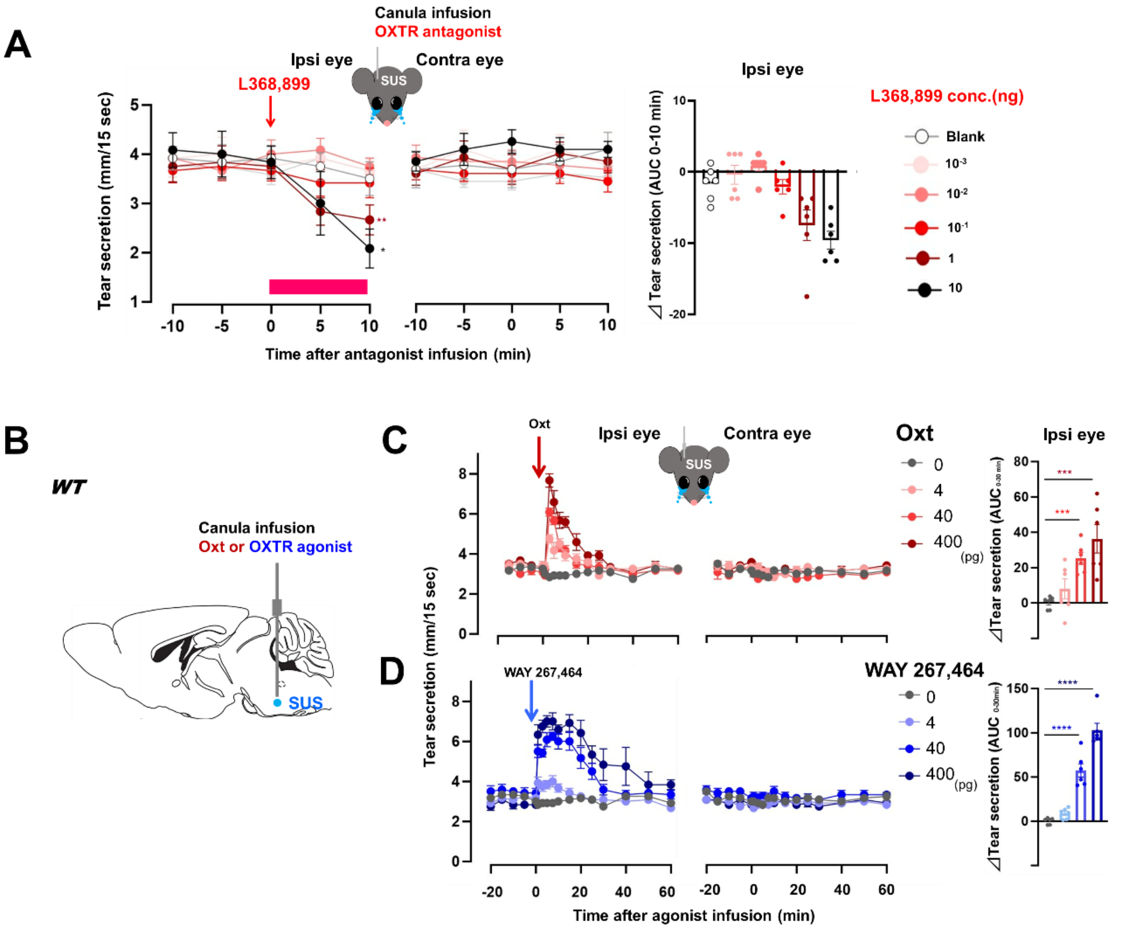


**Figure S9. Changes in tear secretion with pharmacologic activation of SUS OXTR in WT mice.**

A. Effect of pharmacologic suppression of SUS OXTR by OXTR antagonist L368, 899 infusions on basal tear secretion (n = 6 per group).
B. Schematic of pharmacologic activation of SUS OXTR by Oxt or OXTR agonist.
C. Effect of Oxt infusion (n = 6 per group).
D. Effect of OXTR agonist Way 267,464 infusion (n = 6 per group).

****$P < 0.0001$, ***$P < 0.001$. See Table S19 for the statistical analyses.

**Figure S10**

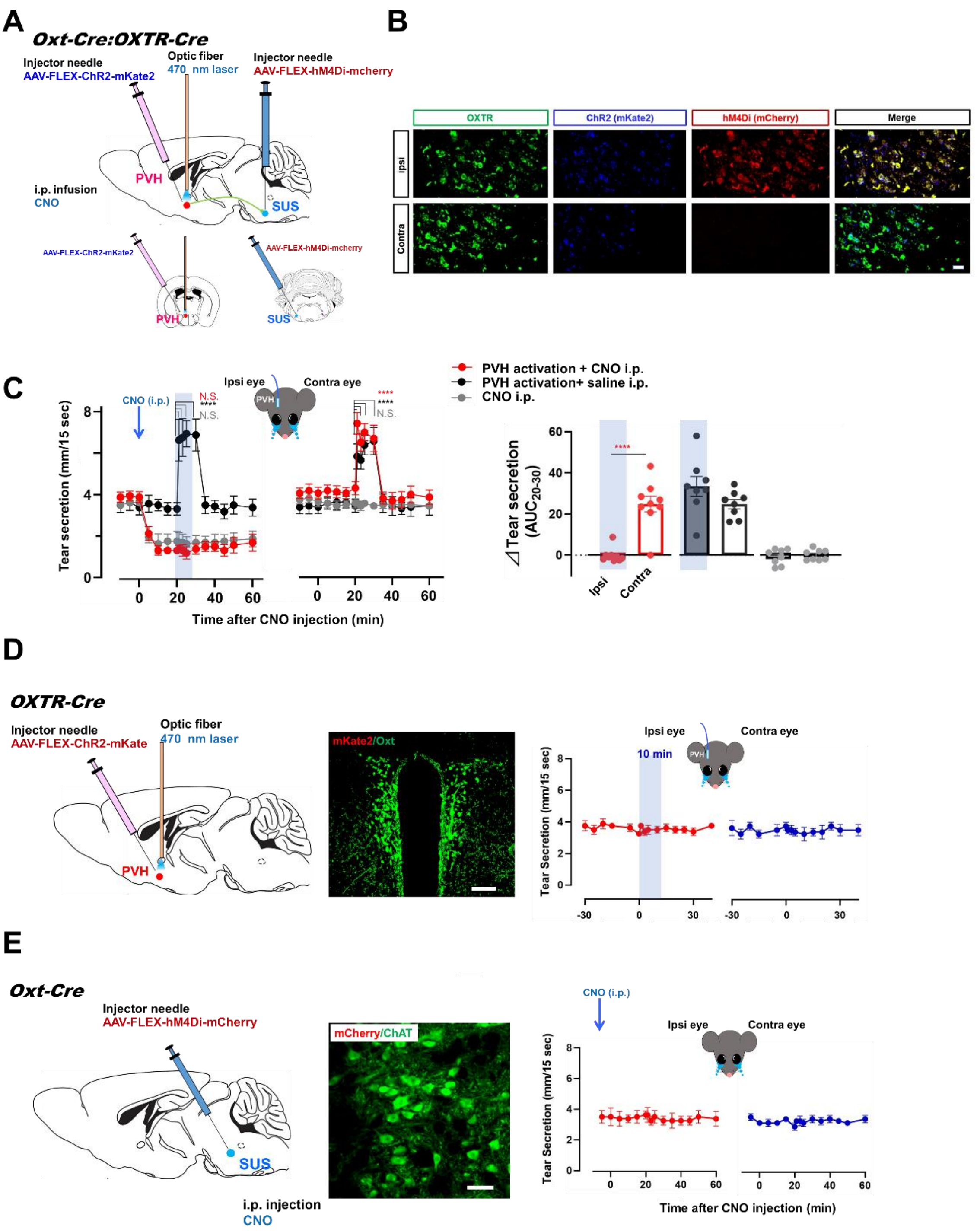


**Figure S10. Functional linkage between SUS-projecting Oxt$^{PVH \rightarrow SUS}$ neurons and OXTR$^{SUS}$neurons in tear secretion,**

A. Schematic of optogenetic activation of Oxt$^{PVH}$ neurons by injection of AAV-FLEX-ChR2-mKate2 and chemogenetic inhibition of OXTR$^{SUS}$ neurons by AAV-FLEX-hM4Di-mCherry in *Oxt*$^{Cre/+}$; *OXTR*$^{Cre/+}$ mice.

B. Histochemical confirmation of ChR2 fused mKate2 and hM4Di fused mCherry in the

SUS. Scale bar represents 50 µm.

C. Changes in tear secretion by activation of $Oxt^{PVH}$ neurons and ipsilateral inhibition of $OXTR^{SUS}$ neurons, dynamics (c, n = 8 for PVH activation + CNO i.p., n = 9 for PVH activation + saline i. p., n = 8 for CNO i.p.) and $\Delta AUC_{(20\text{-}30\ min)}$.

D. Optogenetic activation of PVH OXTR-expressing neurons in $\mathit{OXTR}^{Cre/+}$ mice (n = 4 per group). Schematic of optogenetic activation (left), histochemical confirmation of ChR2 fused mKate2 expression in $Oxt^{PVH}$, changes in tear secretion with optogenetic activation of PVH (right). Scale bar represents 100 µm.

E. Chemogenetic activation of SUS Oxt-expressing neurons in $\mathit{Oxt}^{Cre/+}$ mice (n = 4 per group). Schematic of optogenetic activation (left), histochemical confirmation of hM4Di fused mCherry expression in SUS, changes in tear secretion after activation of SUS (right). Scale bar represents 20 µm.

The shaded areas in the figures represent the periods (C left and D right) or side (C right) of optical stimulation. ****$P < 0.0001$. See Table S20 for the statistical analyses.

**Figure S11**

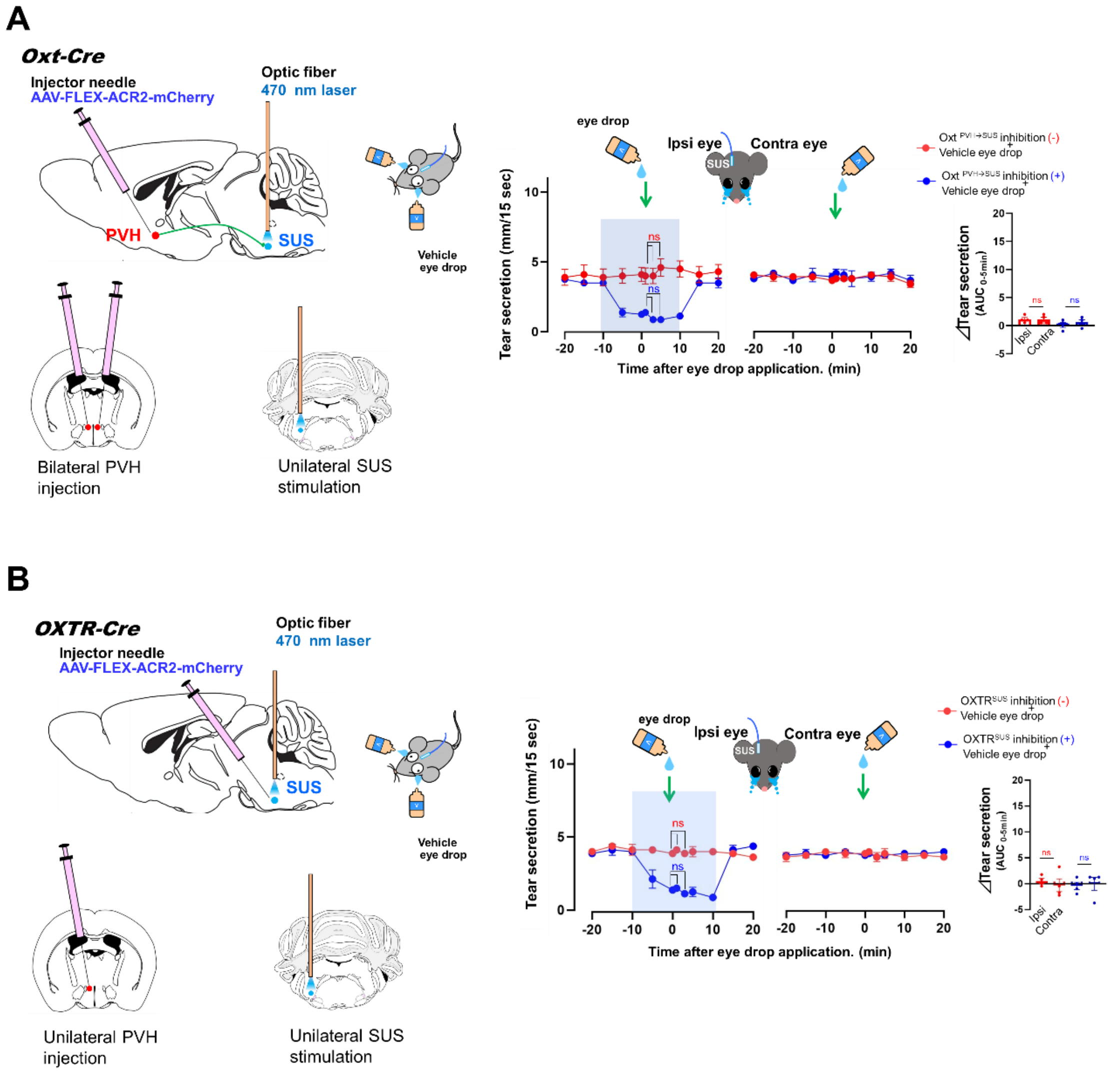


**Figure S11. Effect of vehicle eye drops on tear secretion during optogenetic inhibition of Oxt$^{PVH\rightarrow SUS}$ or OXTR $^{SUS}$ neurons.**

A. Effect of optogenetic inhibition of Oxt$^{PVH\rightarrow SUS}$ neurons (n = 4 per group).

B. Effect of optogenetic inhibition of OXTR$^{SUS}$ neurons (n = 4 per group).

The shaded areas in the line figures represent the periods of optical stimulation. See Table S21 for the statistical analyses.

**Figure S12**

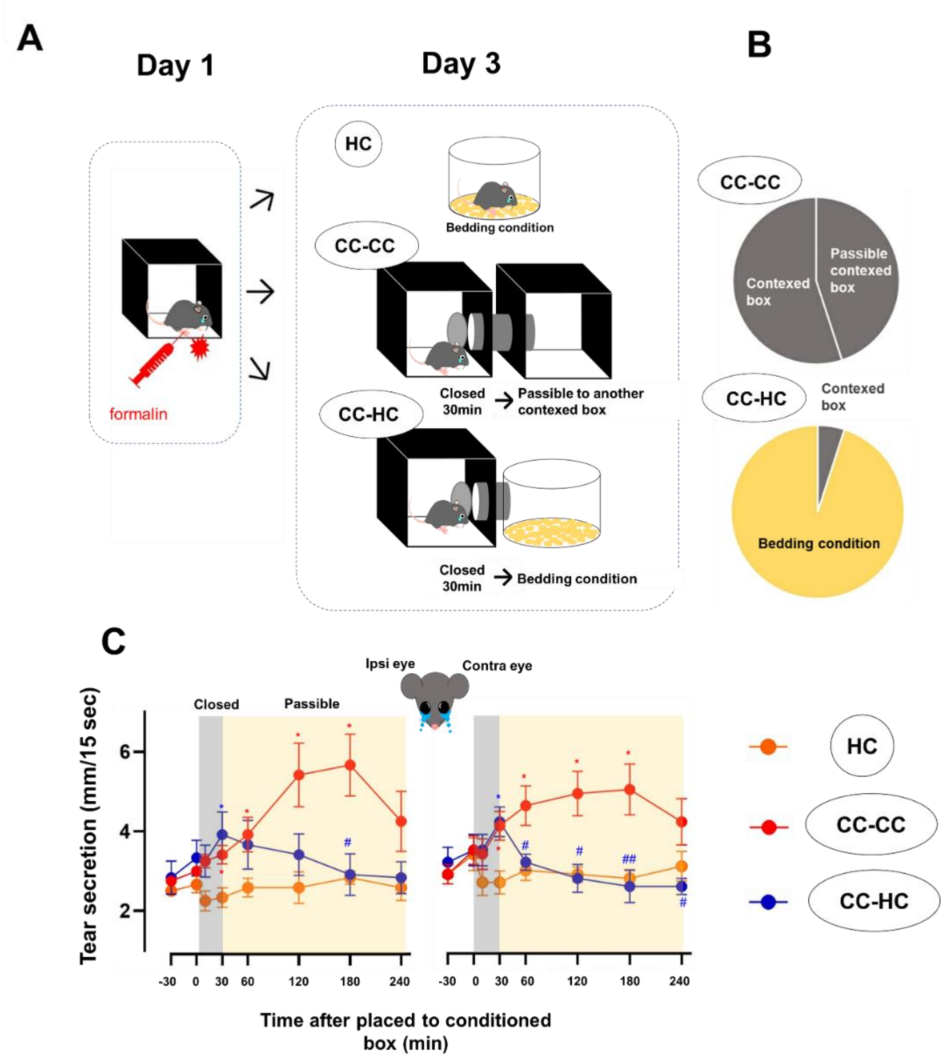


**Figure S12. Confirmation of contextual conditioning-induced tearing.**

A. Experimental schedule.

B. Place preference of the two compartments.

C. Changes in tear secretion in the two compartments (n = 6 per group).

*$P < 0.05$ vs HC, ##$P < 0.01$, #$P < 0.05$ vs CC- CC. See Table S22 for the statistical analyses.

**Figure S13**

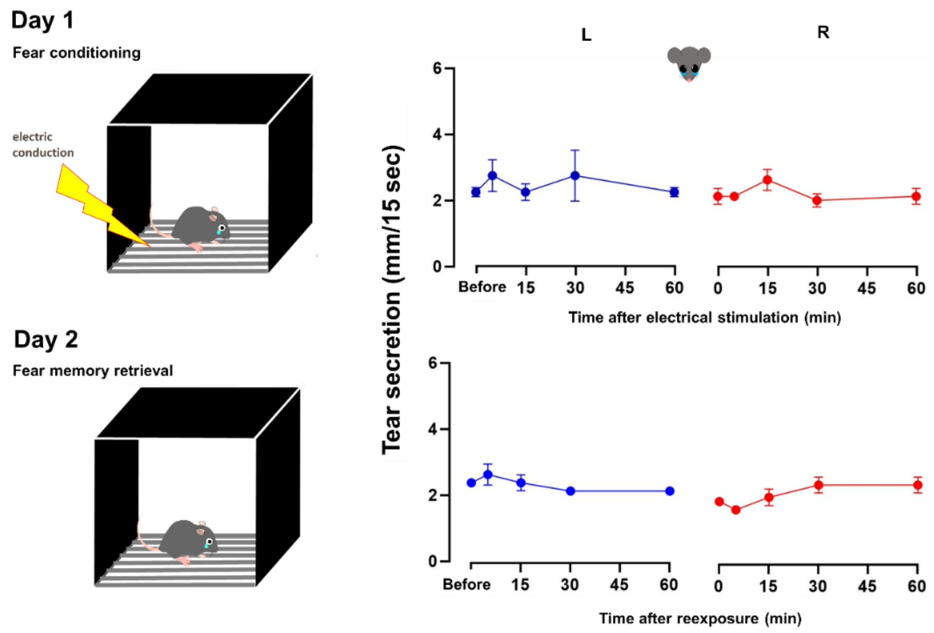


**Figure S13. Changes in tear secretion during contextual fear conditioning by electrical foot shock.**

Contextual fear conditioning by electrical foot shock, nociceptive behavior stimulation that did not evoke tearing (Fig. 6A) did not alter the tear secretion. n = 4 per group. See Table S23 for the statistical analyses.

**Figure S14**

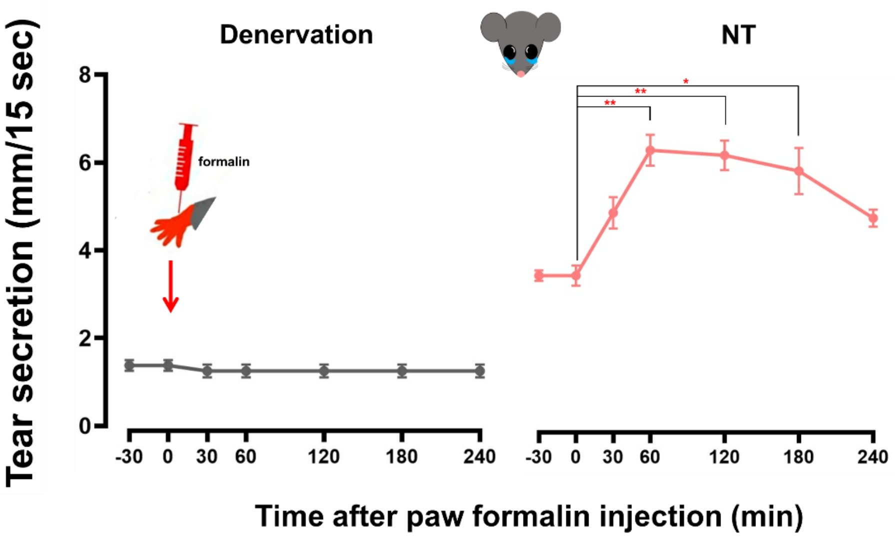


**Figure S14. Changes in tear secretion after hind paw formalin injections under surgical denervation of the lacrimal nerve innervating the LG.**

The lacrimal nerve was surgically denervated unilaterally (n=4 per group). ***P* < 0.01, **P* < 0.05 vs 0 minute. See Table S24 for the statistical analyses.

**Figure S15**

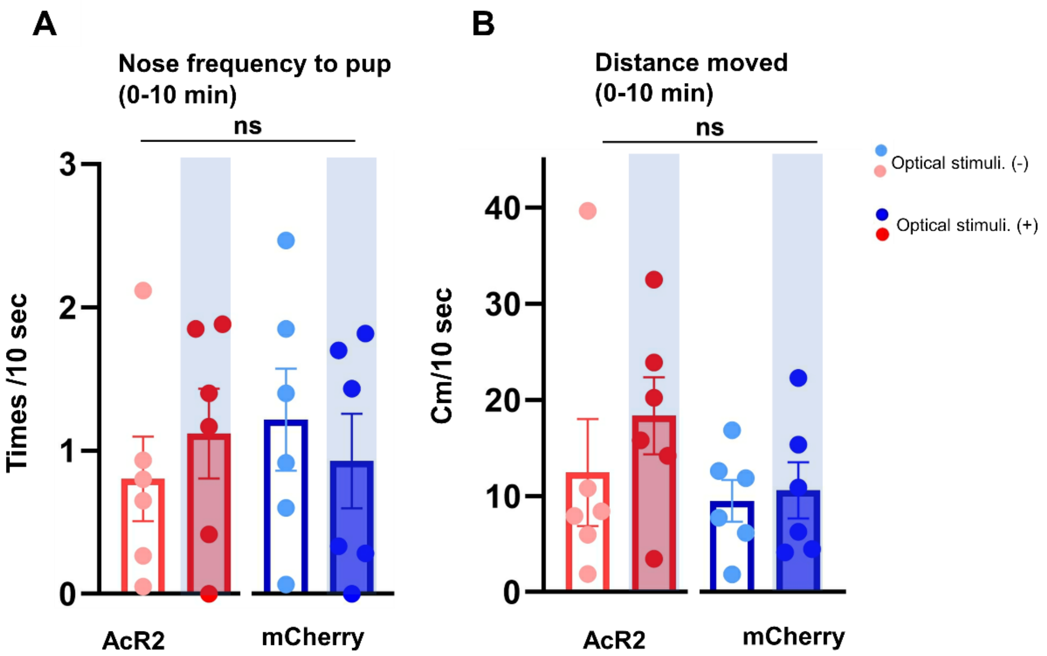


**Figure S15. Changes in mouse pup retrieval behavior with the optical suppression of Oxt$^{PVH \rightarrow SUS}$ neurons. Frequency of nose contact with pups (A), distance moved (B).** mCherry (n = 6 per group), ACR2 (n = 6 per group).

The shaded areas in the figures represent the treatments of optical stimulation. See Table S25 for the statistical analyses.

**Figure S16**

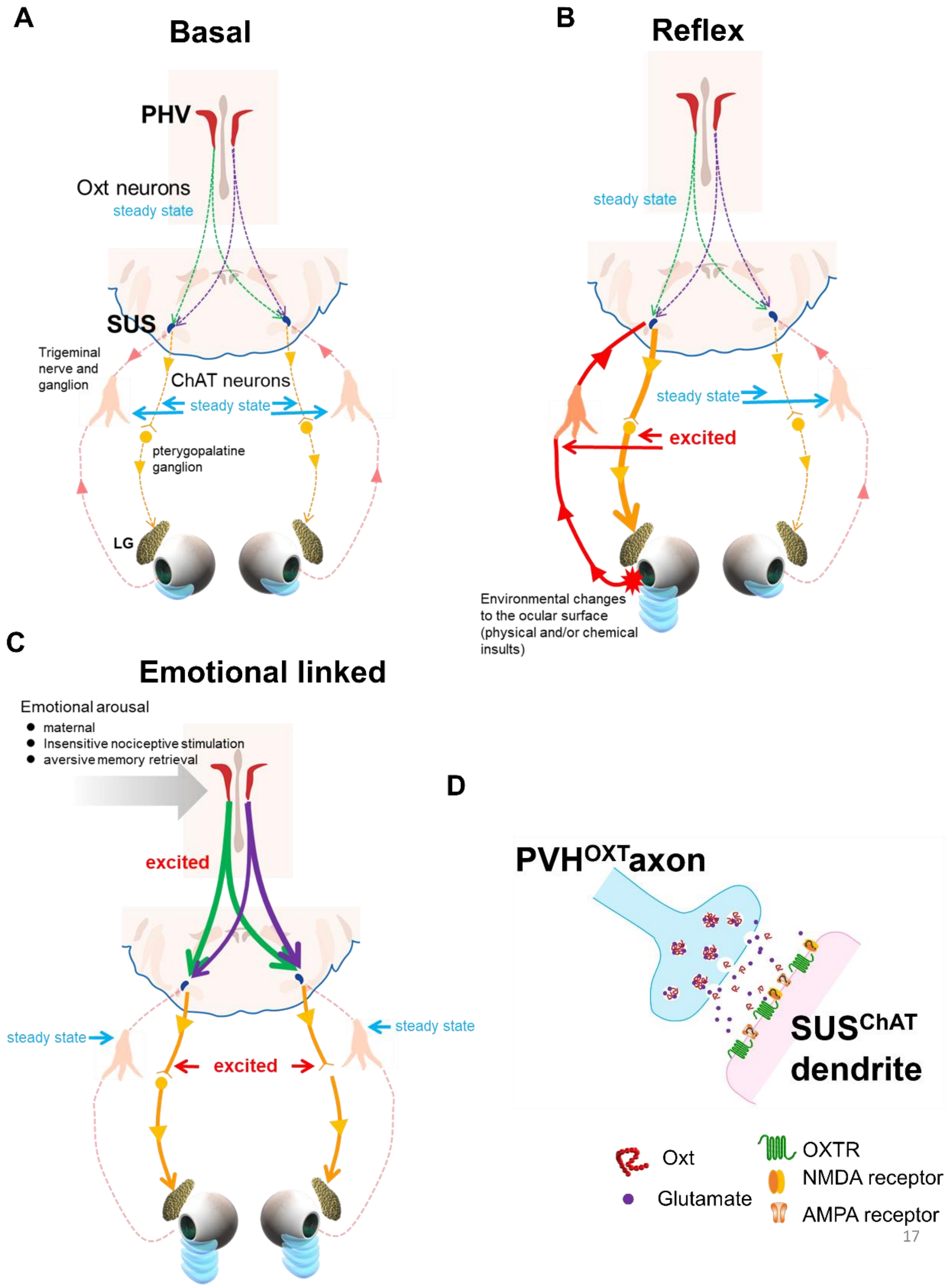


**Figure S16. Schematic diagram of the neural pathways of tearing mediated via the oxytocin system.**

A. **Basal tearing**

B. **Reflex tearing**

C. **Emotional liked tearing**

D. **Oxt released form $OXT^{PVH}$ and $SUS^{OXTR}$ mediates functional connectivity between $OXT^{PVH}$ and $ChAT^{SUS}$ for tearing.**

Tear secretion-related neural response of $Oxt^{PVH}$ is transmitted to the $ChAT^{SUS}$ neurons (Fig. S16A-C) that express $OXTR^{SUS}$. This transmission is modulated by Oxt released from $Oxt^{PVH}$ neurons accompanied by another factor requiring glutamate channel receptor activation (Fig. S16D). The response signals pass through the axons of $ChAT^{SUS}$ and reach the pterygopalatine ganglion (PPG) located behind the posterior maxillary sinus. Postsynaptic fibers from the PPG are then distributed around the LG secretory cells to transmit parasympathetic signals for regulating tear secretion.

**Movies S1A-S1C**

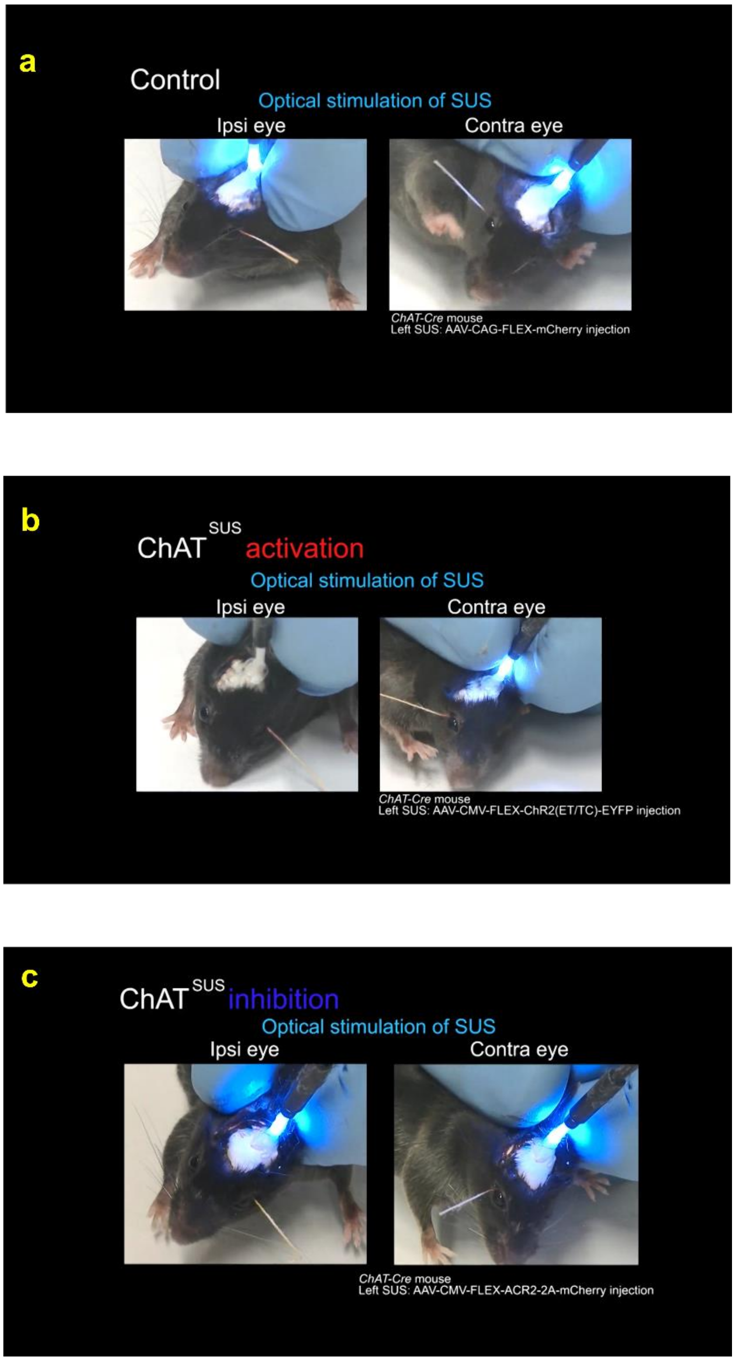


**Movie S2. Change in tear secretion by the optogenetic manipulation of ChAT$^{SUS}$**

A. mCherry (Control)
B. ChR2 (activation)
C. ACR2 (inhibition)

**Movies S2A–S2C**

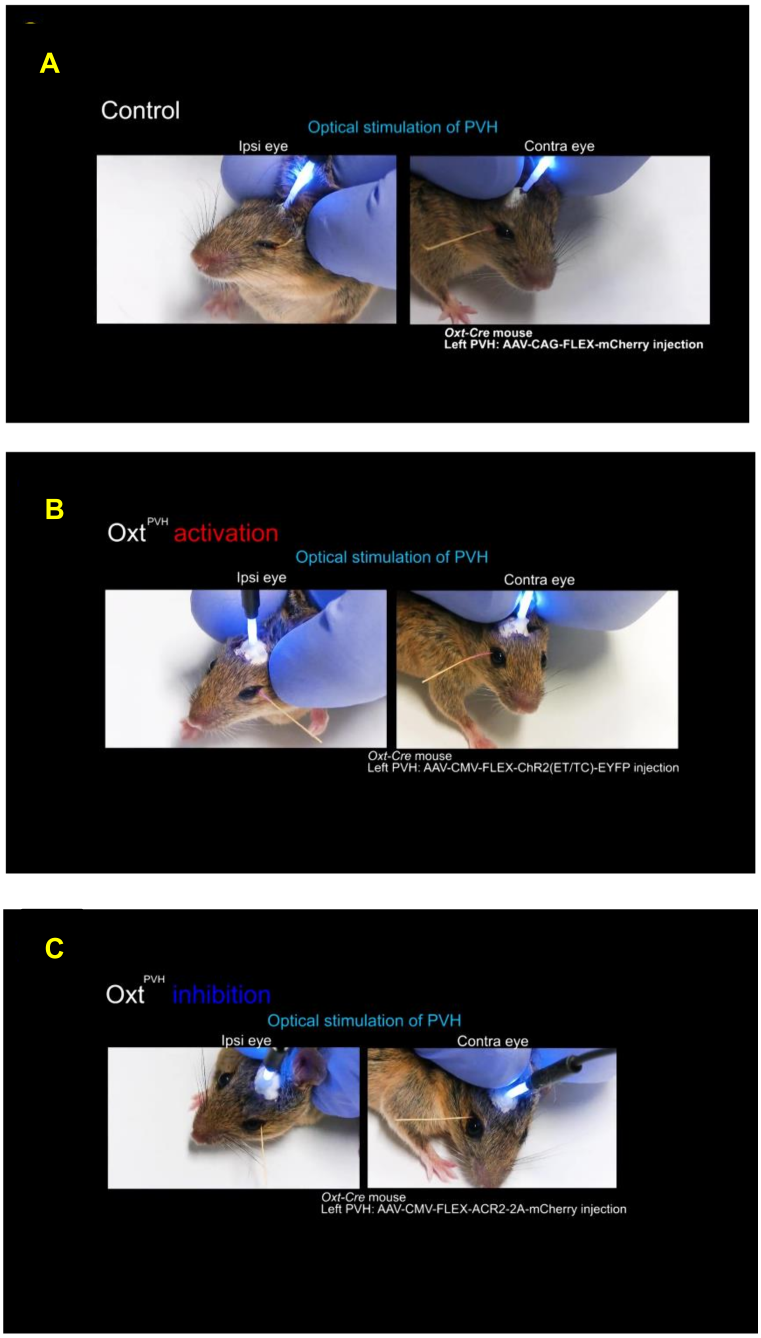


**Movie S2. Change in tear secretion by the optogenetic manipulation of Oxt$^{PVH}$**

D. mCherry (Control)

E. ChR2 (activation)

F. ACR2 (inhibition)

#### Movies S3A–S3B

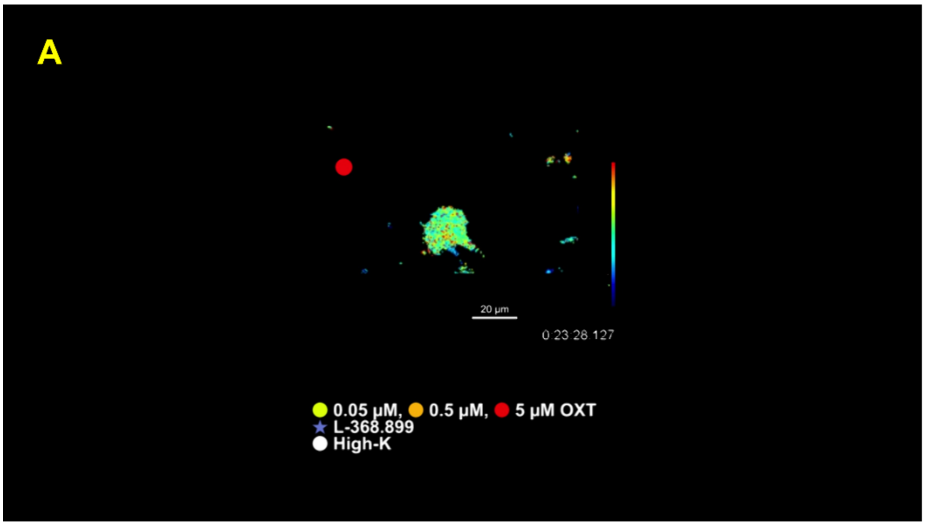


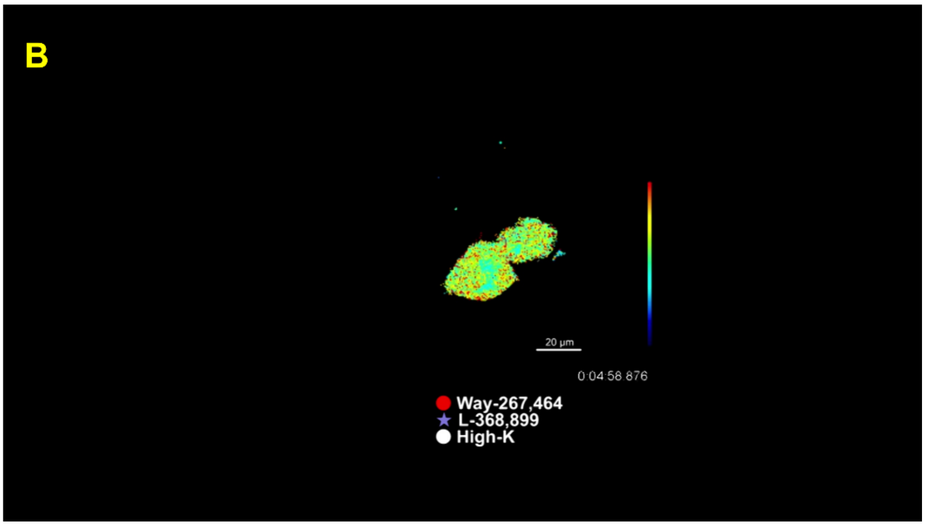


**Movie S3. Dynamics of the FRET ratio in the SUS ChAT neurons stimulated with OXT with/ without OXTR antagonist.**

A. OXT with/ without OXTR antagonist L-368,899

B. OXTR agonist Way-267,464 with/ without OXTR antagonist L-368,899

**Movies S4A-S4C**

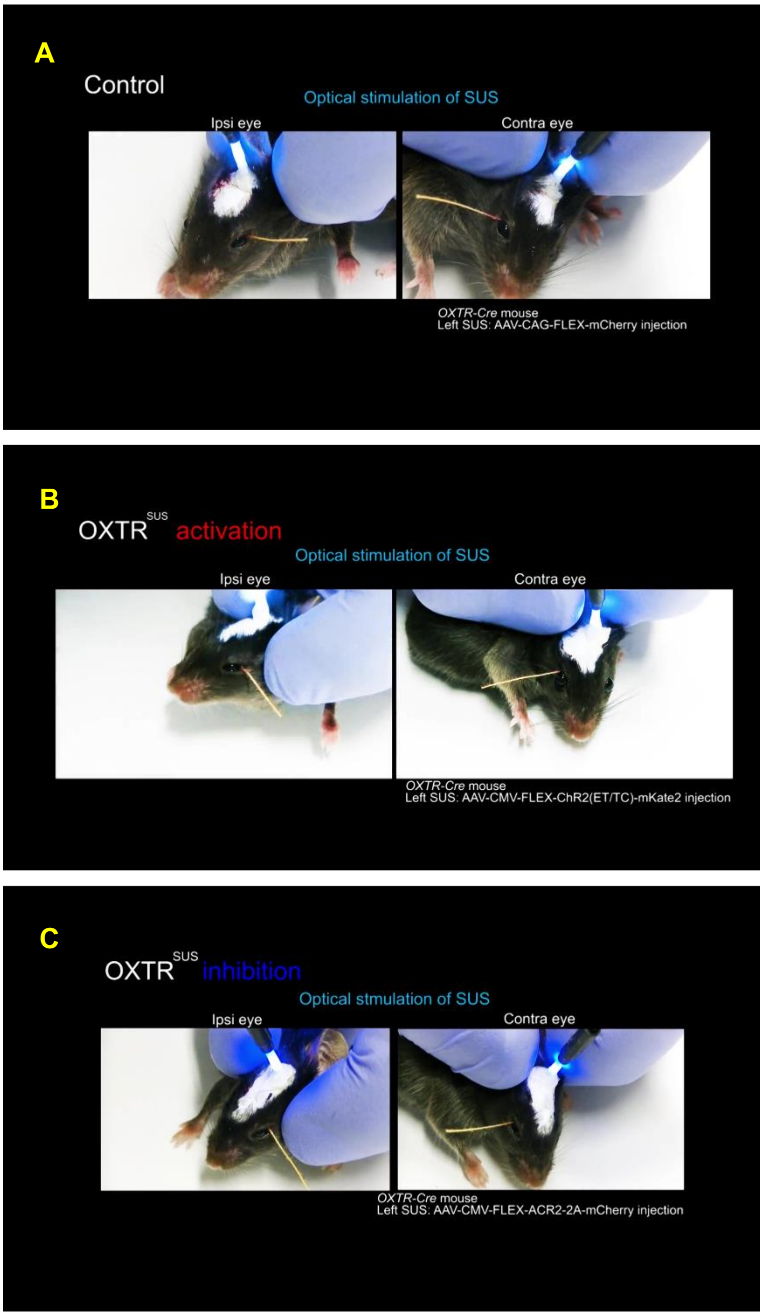


**Movie S4. Change in tear secretion by the optogenetic manipulation of OXTR$^{SUS}$**

A. mCherry (Control)

B. ChR2 (activation)

C. ACR2 (inhibition)

#### References


1. M. Yoshida, Y. Takayanagi, T. Onaka, K. Nishimori, Oxytocin receptor-Venus knock-in mice enable direct visualization of oxytocin receptor-expressing neurons. *Neurosci. Res.* **58**, S222 (2007).
2. P. J. Ryan, S. I. Ross, C. A. Campos, V. A. Derkach, R. D. Palmiter, Oxytocin-receptor-expressing neurons in the parabrachial nucleus regulate fluid intake. *Nat. Neurosci.* **20**, 1722–1733 (2017).
3. K. Jin, T. Imada, R. Hisamura, M. Ito, H. Toriumi, K. F. Tanaka, S. Nakamura, K. Tsubota, Identification of Lacrimal Gland Postganglionic Innervation and Its Regulation of Tear Secretion. *Am. J. Pathol.* **190**, 1068–1079 (2020).
4. F. K. Paxinos G, *Paxinos and Franklin's the Mouse Brain in Stereotaxic Coordinates 3th* (Academic Press Inc., New York, 2007).
5. V. Ferretti, F. Maltese, G. Contarini, M. Nigro, A. Bonavia, H. Huang, V. Gigliucci, G. Morelli, D. Scheggia, F. Managò, G. Castellani, A. Lefevre, L. Cancedda, B. Chini, V. Grinevich, F. Papaleo, Oxytocin Signaling in the Central Amygdala Modulates Emotion Discrimination in Mice. *Curr. Biol.* **29** (2019), doi:10.1016/j.cub.2019.04.070.
6. L. Steru, R. Chermat, B. Thierry, P. Simon, The tail suspension test: A new method for screening antidepressants in mice. *Psychopharmacol. 1985 853*. **85**, 367–370 (1985).
7. S. Tabuchi, T. Tsunematsu, S. W. Black, M. Tominaga, M. Maruyama, K. Takagi, Y. Minokoshi, T. Sakurai, T. S. Kilduff, A. Yamanaka, Conditional ablation of orexin/hypocretin neurons: A new mouse model for the study of narcolepsy and orexin system function. *J. Neurosci.* **34**, 6495–6509 (2014).
